# *Besca*, a single-cell transcriptomics analysis toolkit to accelerate translational research

**DOI:** 10.1101/2020.08.11.245795

**Authors:** Sophia Clara Mädler, Alice Julien-Laferriere, Luis Wyss, Miroslav Phan, Albert S. W. Kang, Eric Ulrich, Roland Schmucki, Jitao David Zhang, Martin Ebeling, Laura Badi, Tony Kam-Thong, Petra C. Schwalie, Klas Hatje

## Abstract

Single-cell RNA sequencing (scRNA-seq) revolutionised our understanding of disease biology and presented the promise of transforming translational research. We developed *Besca*, a toolkit that streamlines scRNA-seq analyses according to current best practices. A standard workflow covers quality control, filtering, and clustering. Two complementary *Besca* modules, utilizing hierarchical cell signatures or supervised machine learning, automate cell annotation and provide harmonised nomenclatures across studies. Subsequently, *Besca* enables estimation of cell type proportions in bulk transcriptomics studies. Using multiple heterogeneous scRNA-seq datasets we show how *Besca* aids acceleration, interoperability, reusability, and interpretability of scRNA-seq data analysis, crucial aspects in translational research and beyond.

## Introduction

Major breakthroughs in our understanding of rare cell types, tissue heterogeneity, cell differentiation and transcriptional regulation have been enabled by the increased resolution in detecting gene expression provided by single-cell RNA-sequencing (scRNA-seq). Encouraged by early successes, pharmaceutical research has also embraced the technology, to accelerate drug discovery. In this context, scRNA-seq is used to better understand disease phenotypes [1], to assess drug targets [2], to characterize microphysiological systems [3] and to measure cell-type-specific pharmacology and toxicity of drug candidates [4], among others. In addition, scRNA-seq assists characterization of *in vitro* and *in vivo* disease and safety models by offering insights in cell-to-cell communication [5], cell activation [6] or differentiation trajectories [7].

Current challenges in the analysis of single-cell transcriptomics data are predominantly related to the biological interpretation of the analysis results rather than to the computation thereof [8]. Whereas the computational part can be automated, biological interpretation requires manual user interaction. By putting our focus on accelerating the cell type annotation process, which is currently a bottleneck in scRNA-seq analyses [9], we aim to streamline the analysis process to ensure that researchers invest their time where it is most effective and to allow for consistent biological investigation. Therefore, we automate and standardize multiple analysis steps as far as possible, in line with current best practices in the community [10–12]. An automated and standardized solution will allow researchers to take full advantage of the rapidly growing amount, size, and scope of single cell data generated [13,14].

Here, we introduce *Besca*, a toolkit for the rapid and standardized analysis of scRNA-seq experiments and the utilisation thereof for the deconvolution of bulk RNA-seq data (Fig. 1). *Besca* is an open-source *Python* library that is compatible with and extends *Scanpy* [15], one of the most established and up-to-date single-cell analysis toolkits. Besides functionalities to analyse scRNA-data, *Besca* also provides the *Besca proportions estimate* (*Bescape*) module, which integrates two cell deconvolution methods: *SCDC* [16] and *MuSiC* [17]. Beyond RNA-focused studies, *Besca* supports analysis of datasets generated by the recently developed CITE-seq (cellular indexing of transcriptomes and epitopes by sequencing) [18] method, hence accounting for multimodal analysis.

**Fig. 1.**
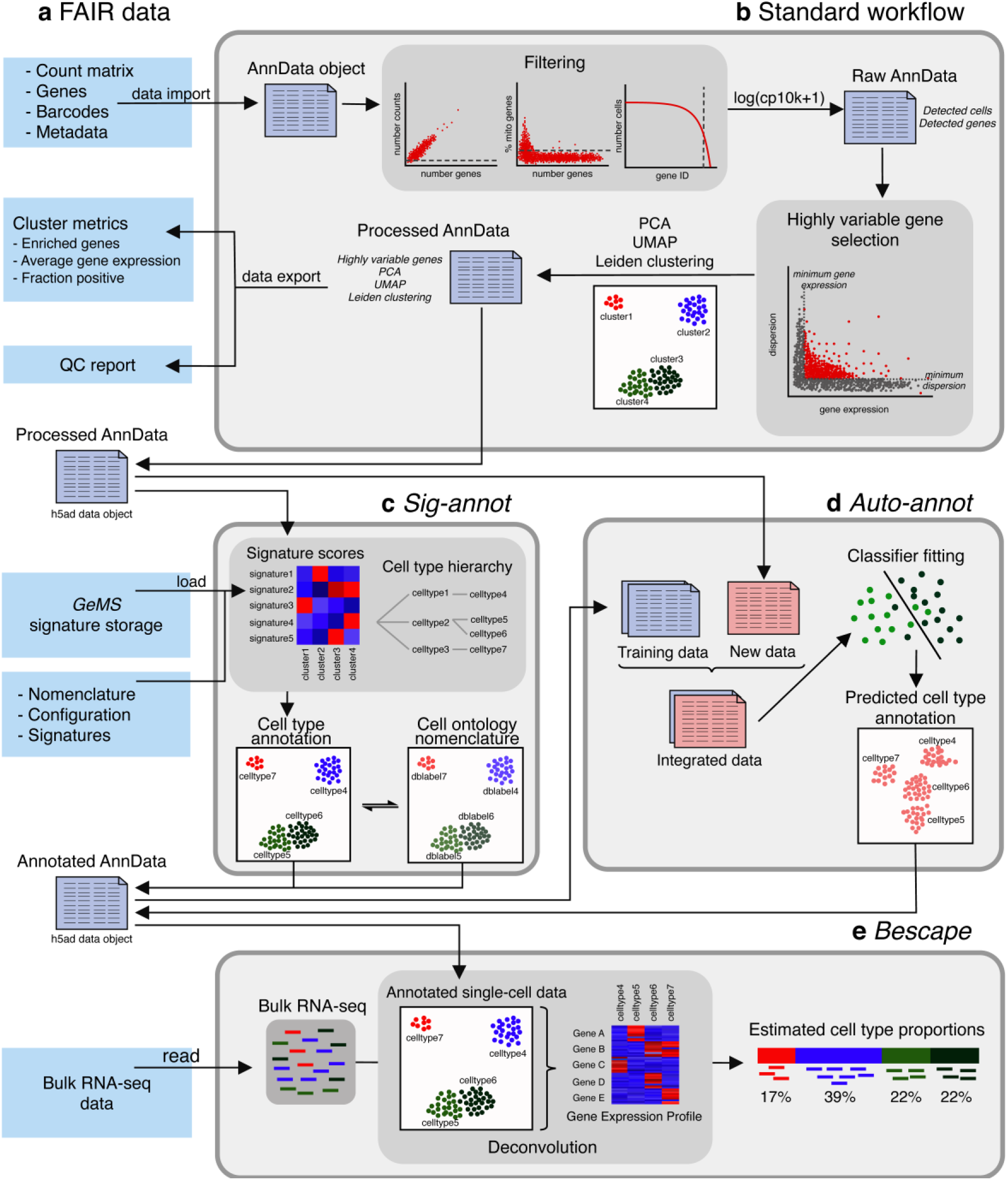
*Besca* provides streamlined single-cell transcriptomics data analysis modules and exchange file formats. **a** Well-defined interoperable input and output file formats, cluster metrics, a quality control report and a signature storage ensure reusability of data. **b** The standard workflow internalizes a raw count matrix and generates a quality control report as well as a processed dataset post filtering, normalization, highly variable gene selection, batch correction, and clustering. **c,d**: Clusters identified from the standard workflow are annotated using either signature-based hierarchical cell annotation (*Sig-annot* module, c) or a supervised machine learning-based algorithm trained on previously annotated datasets (*Auto-annot* module, d). **e** The annotated datasets can be used to deconvolute bulk RNA-seq data based on gene expression profiles generated from annotated single-cell datasets utilizing the *Bescape* module.

As *Besca* builds upon and extends concepts and functions from *Scanpy*, it seamlessly integrates with other ecosystem tools for visualisation or specialised analyses tools such as *scVelo* [19] and *CellRank* (http://cellrank.org/) for cellular trajectory and fate [20] analysis or *Scirpy* [21] for T-cell receptor analysis. We envision *Besca* to accelerate translational research by providing streamlined analysis workflows, ranging from standardized quality control and filtering to harmonised cell annotation.

The *Bescape* module provides a framework to reuse these cell annotations by exploiting scRNA-seq expression profiles for cell deconvolution (Fig. 1e). This adds value to bulk RNA-seq studies, especially in larger clinical settings that do not yet have the capacity to perform scRNA-seq and where signals are often confounded by heterogeneity related to distinct cell type composition [22]. The resulting estimated cell compositions can then be used directly as biomarkers or as covariates towards getting more robust differential gene expression results for understanding disease biology or treatment responses.

In this manuscript, we exemplify how using *Besca* makes analysis results more comparable between studies. We also demonstrate how to transfer learnings from one study to another, for instance by reusing cell type annotations, and from one application to another, for instance by using single-cell gene expression profiles for cell deconvolution of bulk RNA-seq. To demonstrate how *Besca* can be applied to a wide variety of biological samples, we reprocessed publicly available single-cell data from ten studies (see Table 1 and Methods). We show how the *Besca* toolkit can be used to obtain biological insights quickly and generate reusable results from these highly diverse datasets. Further examples can be found in the supplementary material, example workbooks on GitHub (https://github.com/bedapub/besca_publication_results), and in the tutorials available from the documentation (https://bedapub.github.io/besca/).

**Table 1.**
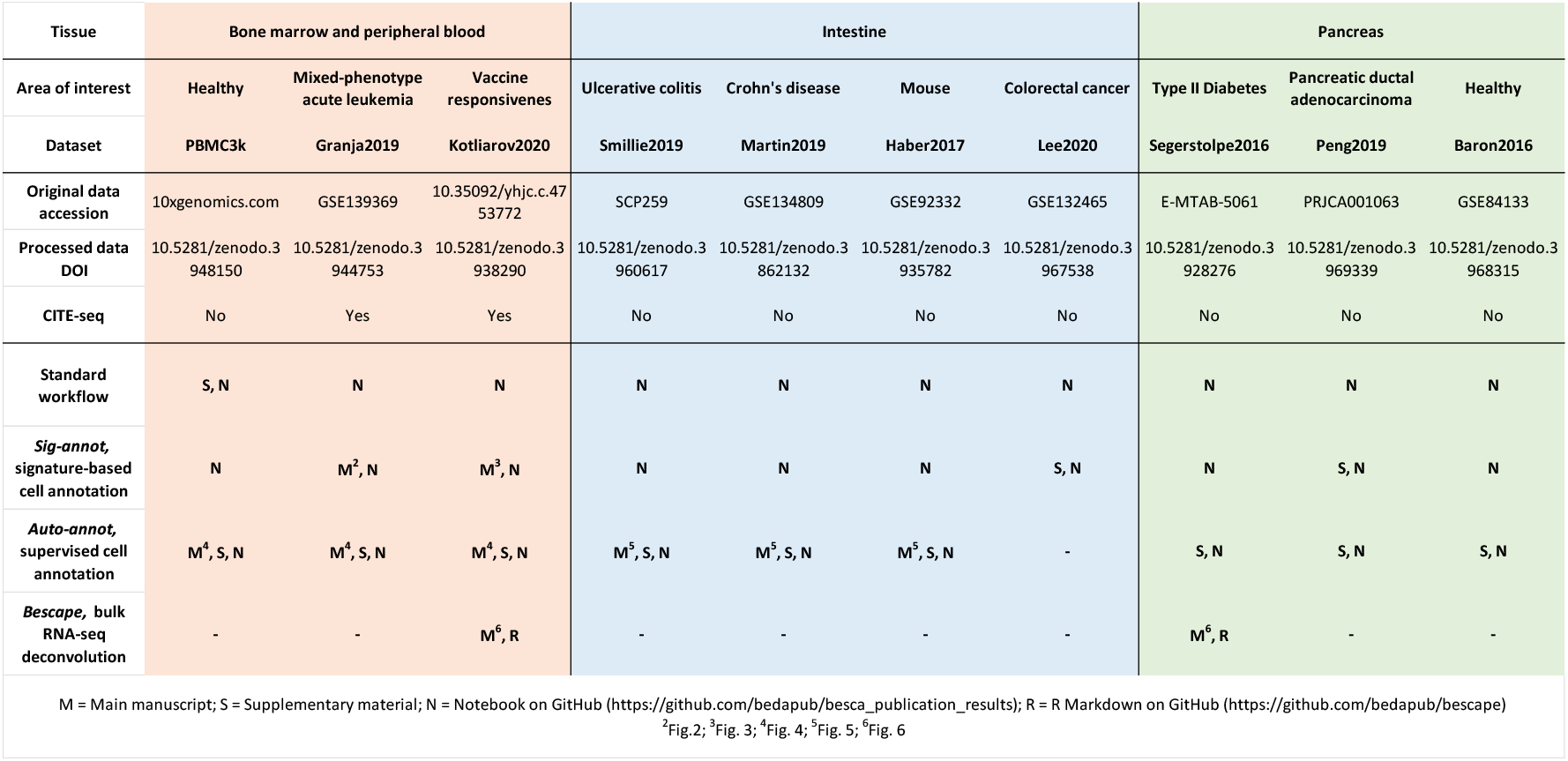
Dataset overview, including hematopoietic cells of peripheral blood and bone marrow (orange), intestine (blue) and pancreas (green) in health and disease.

## Results

### A standard workflow streamlining scRNA-seq and CITE-seq analyses

The *Besca* standard workflow offers a standardized series of steps, starting from a gene-by-cell count matrix and ending with cell clustering (Fig.1b). Based on *Scanpy* [15], the workflow provides standard processes to treat single-cell transcriptomics data in a reproducible and comparable manner. Good practices and FAIR (findability, accessibility, interoperability, reusability) principles [23] enable comparisons between all datasets analysed with *Besca* improving translational research. The steps of single-cell analysis are described at length by Luecken and Theis [11]. *Besca*’s standard workflow detailed in the Methods follows these steps and in addition allows for the processing of CITE-seq [18] data.

The standard workflow generates a quality control (QC) report and a log file which summarize the performed analysis (Fig. 1a). For future reuse, all of the analysis results are written to files in interoperable data formats (see Methods) including output files of precomputed metrics, such as average gene expression or marker gene rankings (Fig. 1a). Additional downstream analyses such as automated cell type annotation can be run directly on the output of the standard workflow. The cell type annotation of the clusters can be performed using the *Sig-annot* (Fig. 1c) or *Auto-annot* (Fig. 1d) methods described thereafter and a re-clustering framework is available to decipher cell populations with higher resolution. In addition, functions for recurrent visualizations are implemented to illustrate gene expression variation under certain conditions (e.g. treatment effect) or to show the cell type composition found in the analysed dataset. The standard workflow and subsequent manual cell type annotation are exemplified in Supplementary Figure S1 utilizing the PBMC3k dataset.

### A gene signature management system

The integration of multiple scRNA-seq datasets allows for the accumulation of knowledge and insights about biological tissues, cells, cell states, and diseases. As the development of suitable scRNA-seq integration data increases, a key challenge in single-cell data analysis workflows is the accurate dissemination of this knowledge and the appropriate reuse of the information gathered. In particular, it is of utmost importance to be able to re-apply gene signatures extracted from individual studies across studies and within analyses. To this end, we connected *Besca* to the *Geneset Management System* (*GeMS*) (https://github.com/bedapub/GeMS). *GeMS* is a light web-based platform that enables the centralized management of genesets using structured formats and a local application programming interface for geneset information retrieval and organization. The application is built on top of the *Flask* micro-framework (https://flask.palletsprojects.com) using *MongoDB* (http://www.monogdb.com), an open-source, document-based database as its backend.

Once *GeMS* is deployed, *Besca* allows the export of gene signatures to the *GeMS* database (for example a geneset of marker genes from distinct populations) and the retrieval of user-defined signatures (Fig. 1a). It is also possible to check for geneset similarity to avoid redundancy within the database and check for signature specificity. *GeMS* is distributed with initial public genesets extracted from *Reactome* [24], *CREEDS* [25], *CellMarker* [26] and *MSigDB* [27,28] and can be filled with new genesets. *Besca* allows for direct usage of these genesets for signature enrichment analysis and can compute bi-directional scores combining up and down-regulated genes into one metric. *Besca* is distributed with signatures related to different tissue types including hematopoietic, intestinal and pancreatic cell types as well as an extended list of immune-related signatures.

### Automated and harmonized cell type annotation

Cell type identification in scRNA-seq poses great challenges, mainly related to the lack of a biological consensus of what a cell type actually represents and a patchy overview of existing cell types and their identity footprints on the transcriptomic level [29,8]. During recent years, a large number of approaches and computational methods have been developed to address the attribution of cells to discrete types, however a one-fit-for-all approach is still lacking [30]. At the most basic level, cell types are attributed iteratively to individual clusters after manual inspection of the expression of a handful of markers according to expert biological knowledge of the studied system. Importantly, the vast majority of scRNA-seq-based publications have taken this approach in the past (see e.g. [31–39]). However, such an approach is limited by the availability of expert knowledge, does not scale to processing a large number of samples, and is poorly reproducible across individual studies.

In order to standardize this process, while maintaining the flexibility of adjusting marker genes and expression cut-offs across studies according to prior knowledge, we developed *Sig-annot* (Fig. 1c), a *Besca* module that provides a hierarchical signature-enrichment approach for cell type annotation (see next paragraph). To guarantee consistency across studies and communities, beyond scRNA-seq, the proposed cell type annotation schemas are based on the *Cell Ontology* [40], which is accessible via the *Experimental Factor Ontology* [41]. The controlled vocabularies at different cell type hierarchies are summarized in an annotation sheet (Supplementary Table S1) and can be easily extended with further cell types. Newly generated cell type annotations in this manuscript provide the most fine-grained annotation as DBlabel assignment, which follows the *Cell Ontology* whenever possible, as well as higher level annotations according to the annotation sheet.

### Sig-annot, Besca’s signature-based hierarchical cell annotation schema

*Sig-annot* is *Besca*’s streamlined version of the manual process of cluster attribution based on marker gene enrichment including ready-to-go annotation schemas for a broad range of cell types, with a particular focus on immune cells. The flexible, multi-level identification schemas are based on a configuration file containing the cell types and their relations as well as the corresponding cell type signatures (see Methods). Default configuration files for human and mouse are provided, covering a large range of tissues and cell types (human: Supplementary Table S2, mouse: Supplementary Table S3). These files are easily customisable and users are free to provide additional schemas or annotations. The corresponding cell type signatures provided with *Besca* (Supplementary Table S4) are derived and adapted from various scRNA-seq experiments and publications, with subsequent manual curation. As demonstrated here, they can be applied across tissues and potentially even species (with some dataset-specific adjustments) and represent a fast and consistent way of determining the most likely cell type composition in complex, large-scale scRNA-seq experiments.

For convenience, we have implemented various functions to guide the annotation based on the *Sig-annot* framework, and also provide visualisation at individual steps. For instance, one can visualise the relation between the individual cell types as a graph (Fig. 2a), plot the enrichment of individual signatures across all clusters in the dataset as a heatmap (Fig. 2b), directly generate annotations at distinct levels in the cell hierarchy and add these in bulk to the *AnnData* (https://anndata.readthedocs.io) metadata.

**Fig. 2.**
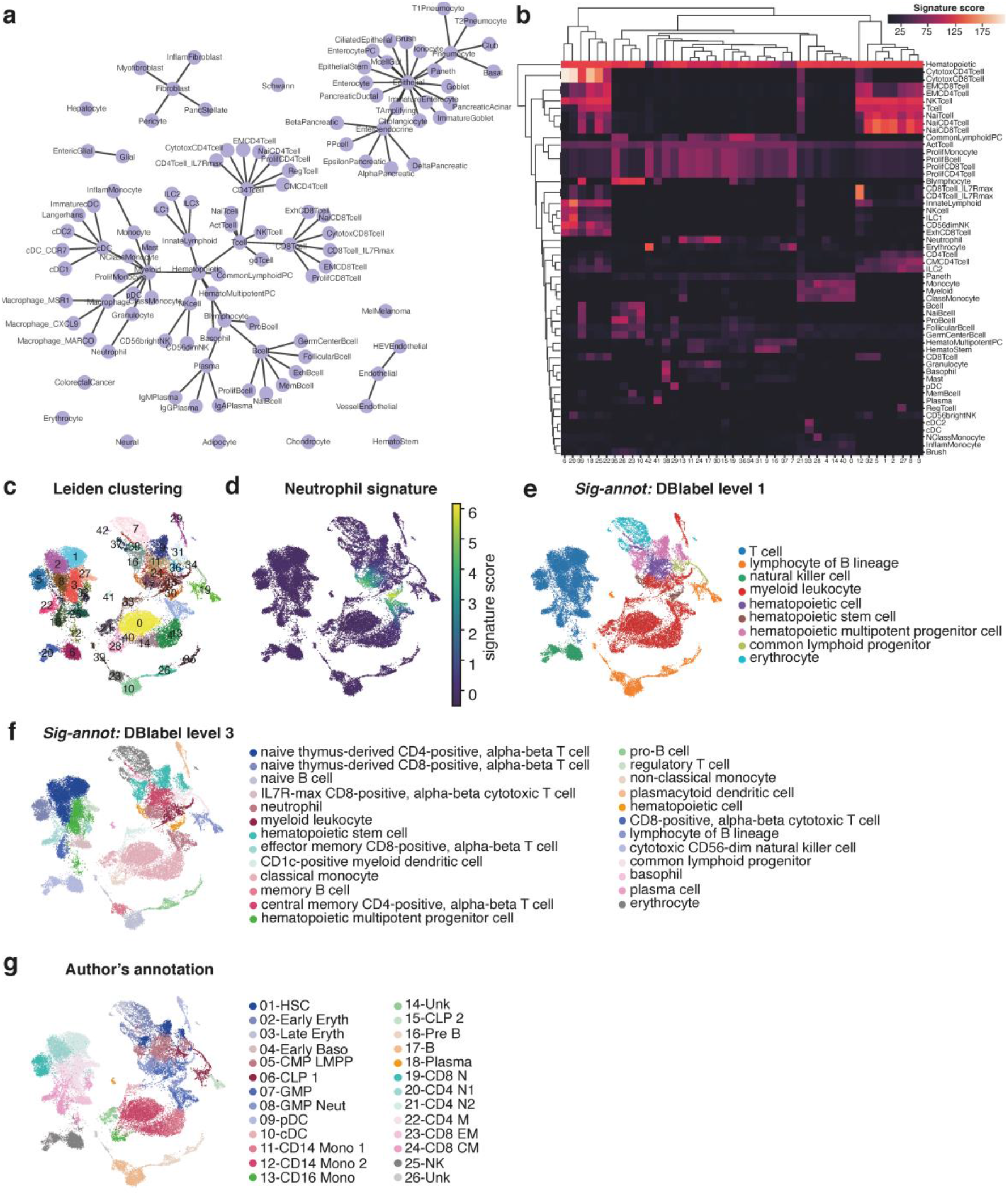
*Besca*’s *Sig-annot* module applied. **a** Overview of the cell type hierarchy provided with *Besca*’s *Sig-annot* module and employed for annotating the datasets in the current manuscript. **b-e**: Granja2019 data containing hematopoietic cells of multiple healthy donors from blood and bone marrow, probed by CITE-seq. **b** Hierarchically clustered heatmap showing enrichment of main signatures employed in the annotation across Leiden clusters, facilitating the evaluation of cluster attribution. **c** Overview of clustering in 2D UMAP space. **d** Overview of one of the signatures employed in cell annotation; neutrophils are typically rare in scRNA-seq experiments because of their sensitivity to cell isolation protocols, but can be clearly detected in the Granja2019 dataset based on the *Besca* included signature. **e** *Sig-annot* cell type attribution at level 1, consisting of major cell types such as T cells and myeloid cells. All detected populations are broadly consistent with the original annotation (g). **f** *Sig-annot* cell type attribution at level 3, the highest resolution provided in *Besca*’s cell annotation schema. The detected populations are consistent with the original Granja annotation (g), cover T cell subsets with higher granularity and attribute the previously unknown (‘14_Unk’ and ‘26_Unk’) clusters as well. **g** Original cell type attribution as obtained from Granja *et al*. Annotated cell populations are highly consistent with clusters obtained from the reanalysis of the original data following the *Besca* standard workflow.

To exemplify this approach and its utility across samples of various origin and characteristics, we apply it to recent publicly available datasets covering most known hematopoietic cell types [42] and show that we are able to reproduce and enhance the original expert-driven annotations [31,32] (Fig. 2 and 3). As one of the datasets also contains information on the expression of a large number of surface protein markers, we can confirm that our cell type attribution is in line with our current protein-level understanding of hematopoietic cell biology (Fig. 3b and d, Supplementary Figure S2).

**Fig. 3.**
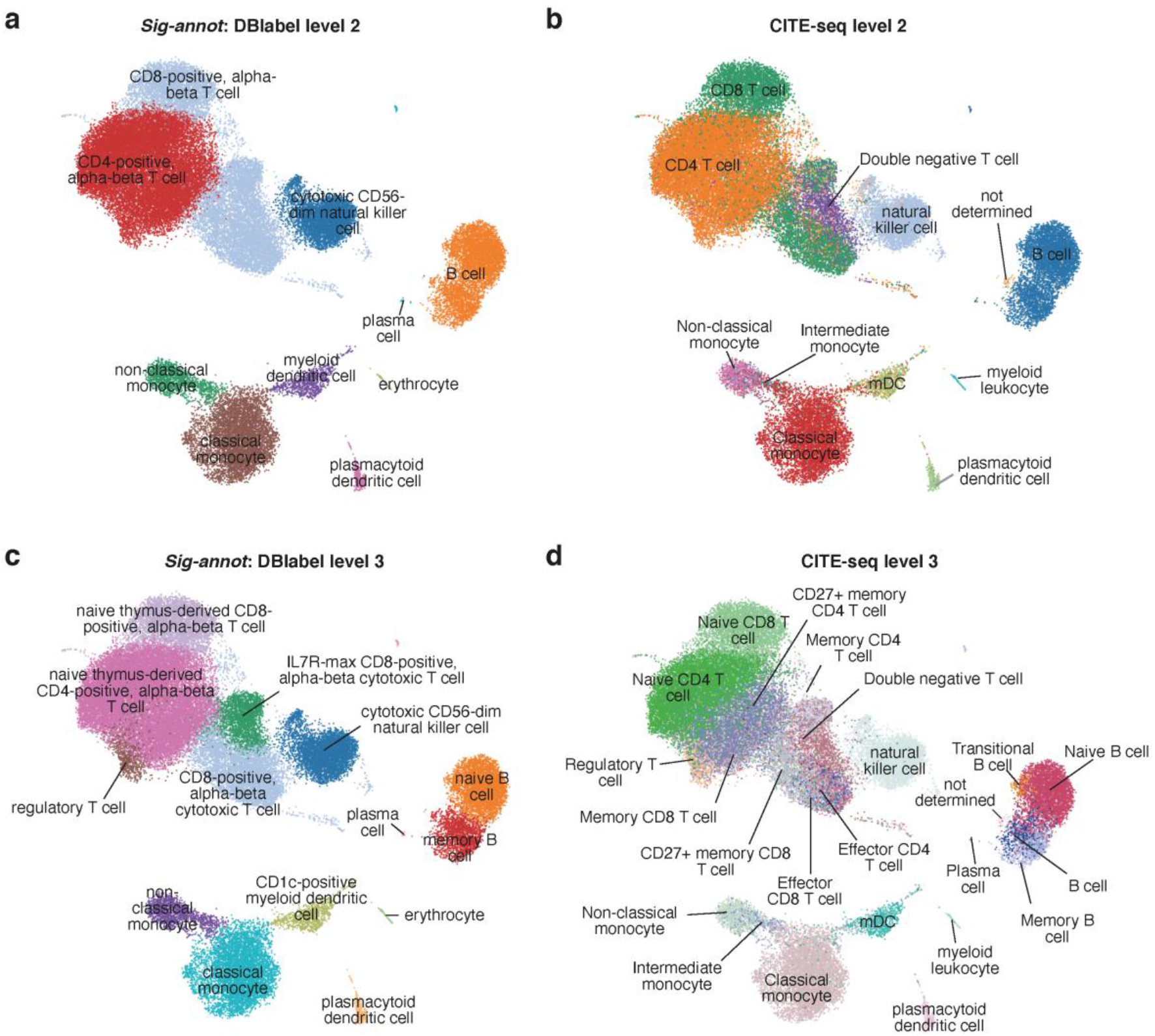
*Sig-annot* applied to Kotliarov2020 data containing hematopoietic cells of multiple healthy donors from blood, probed by CITE-seq. **a** RNA signature-based cell type attribution at level 2, consisting of cell subtypes such as CD4+ T cells and classical monocytes. **b** Protein-marker based annotation using a gating method of classical FACS markers at a similar hierarchical depth as described in (a). Cell attribution is highly consistent with the automated RNA based results. **c,d** RNA signature-based (c) and protein-based (d) cell type attribution at the most fine-grained level 3. Even immune cell subtypes such as memory versus naive B cells or rare populations such as regulatory T cells and plasmacytoid dendritic cells are correctly attributed.

We note that in our annotations, we employed the same set of signatures and configuration files, successfully obtaining consistent annotations of hematopoietic cells derived from independent experiments, each with distinct levels of resolution and cell type frequency and representation, covering human blood and bone marrow. Importantly, our approach is automated, in the sense that only minimal changes (if any) are required for re-annotating each dataset should e.g. filtering/clustering be modified. It is also fully reproducible if the signature matrix and configuration files are stored for each annotation event. The distinct levels provide flexibility in terms of the annotation depth - one can easily choose to inspect differences between myeloid cells and T cells, or alternatively examine myeloid cell subsets, as each cell is attributed all hierarchical annotation levels present in the configuration file. Finally, we demonstrate that our approach is also applicable to more complex settings such as heterogeneous tumor samples, as exemplified by the annotation of publicly available colorectal cancer and pancreatic cancer data (see Supplementary Figures S3 and S4).

### *Auto-annot*, *Besca*’s supervised machine learning module for cell type annotation

In addition to the signature-based annotation approach, *Besca* provides the *Auto-annot* module (Fig. 1d), a supervised machine learning workflow for automated cell type annotation based on well annotated training datasets. Recently, supervised machine learning has become a popular alternative to signature-based cell type annotation [43–46]. Benchmarking studies of such methods revealed that tailored single-cell classifiers or deep learning algorithms do not perform significantly better than conventional general purpose machine-learning methods [30,47]. Therefore, we implemented methods for supervised machine learning based on support vector machines (SVMs) or logistic regression. One or multiple annotated reference datasets can be used to train a classifier for the annotation of a test dataset. Further details of the implementation are described in the Methods.

We demonstrate the application of *Auto-annot* on scRNA-seq data from healthy PBMCs. The datasets Kotliarov2020 and Granja2019 (Table 1, [31,32]) were used to train a logistic regression model, which was then tested on the PBMC3k dataset (Table 1, https://www.10xgenomics.com/). The training data includes far more cells and is annotated more fine-grained, a scenario we expect when training on deeply annotated datasets derived from cell atlases and predicting cell identities in smaller newly sequenced datasets.

The resulting automated annotation (Fig. 4b) broadly reproduces the reference annotation (Fig. 4a,f), and also highly overlaps with the unsupervised Leiden clustering from *Besca*’s standard workflow (Fig. 4c). For B cells, it provides even higher resolution than the reference annotation correctly separating them into memory and naive B cells (Fig. 4b,f), as independently confirmed by the according signatures (Fig. 4d,e). The automated annotation for T cells shows some ambiguity, which reveals the limitations of the method (Fig. 4a,f). Still, the specific IL7R-max CD8 T cells were correctly identified (Fig. 4f) showing that accurate subdivisions within T cells are possible. In order to avoid false positive annotations it is possible to set a threshold for cells with low annotation scores. The threshold approach labels most of the ambiguous T cells as unknowns (Fig. 4g and Supplementary Figure S5), removing almost all misclassifications at the cost of some cell types. As a result, central memory CD4 T cells remain virtually undetected. However, little changes occur when it comes to other cell types, including IL7R-max CD8 T cells, suggesting that this approach indeed only flags out ambiguous attributions.

**Fig. 4.**
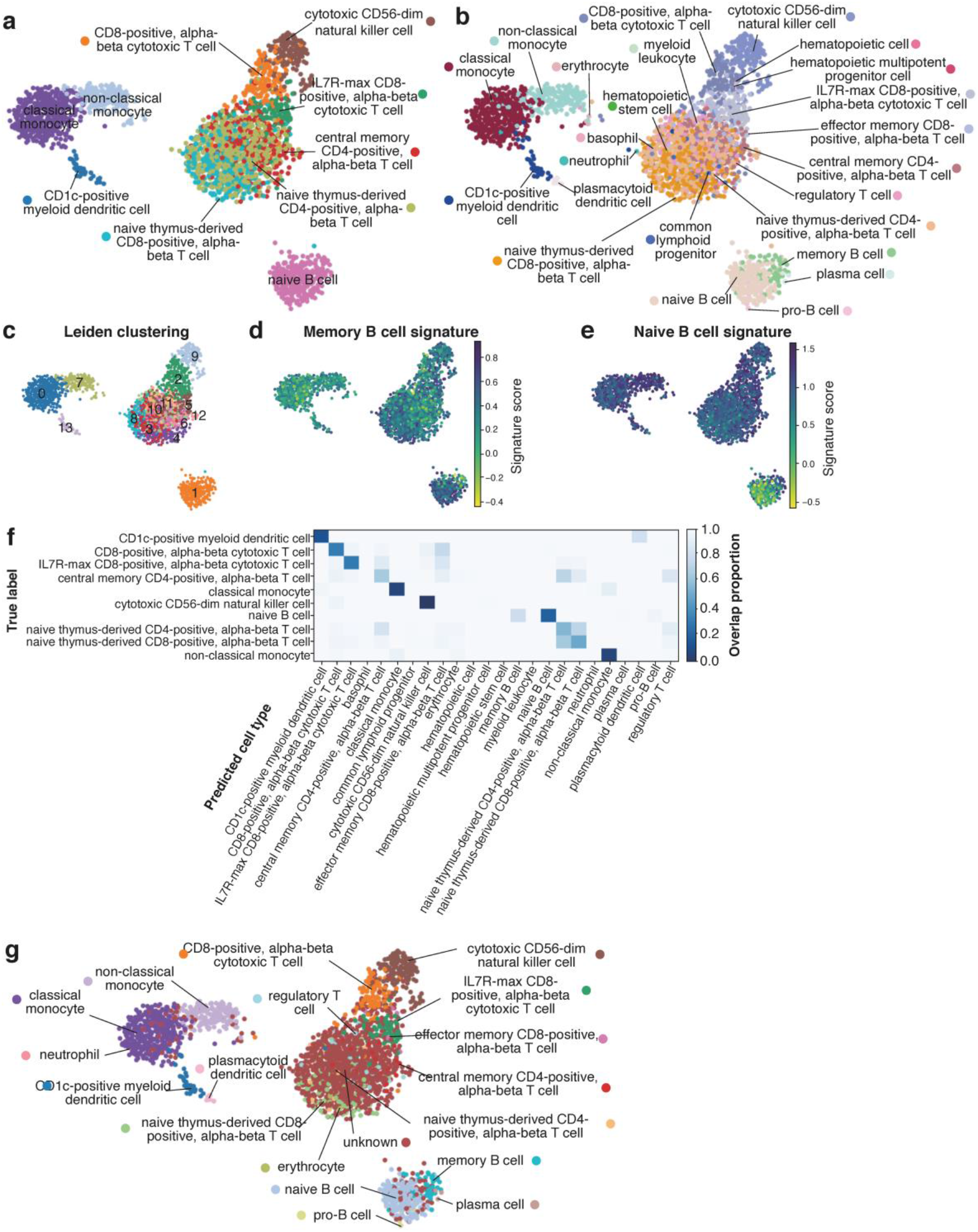
*Auto-annot* applied to PBMCs using a logistic regression model trained on the Kotliarov2020 and Granja2019 datasets and tested in the PBMC3k dataset. **a** Overview of DBlabel annotations in 2D UMAP space for the PBMC3k test dataset. **b** *Auto-annot* largely recovers the original cell types. Finer divisions are uncovered in B cells, but resolution is lost for some T cell subtypes. **c** Overview of Leiden clustering in 2D UMAP space shows high overlap with predictions and illustrates the difficulty of finding subclusters in overlapping T cell communities. **d** The memory B cell signature supports the separation of the B cell cluster in (b). **e** Idem for the naive B cell signature. **f** The confusion matrix shows that misclassifications, if they do occur, generally misannotate very similar cell types. **g** Overview of *Auto-annot* labels with threshold. Ambiguity in some T cell subtypes leads to classification as unknown, all other cell types remain identified. The corresponding confusion matrix can be found in Supplementary Figure S5.

It is notable how accurate the supervised approach works with a fine-grained training annotation. Still, an automated annotation based on less fine-grained cell types leads to even clearer results in the sense that multiple different cell types being co-located in the same broad cell type class from the reference annotation does not occur when we applied it to broader cell types (see Optimised Classes in the Supplementary Material and Supplementary Figure S6).

We performed additional cross-validation of the supervised *Auto-annot* approach on hematopoietic cells using the Granja2019 and Kotliarov2020 datasets on their own (see Supplementary Figures S7 and S8) and on pancreatic cells utilizing the Segerstolpe2016, Peng2019, and Baron2016 annotations in three different combinations (see Supplementary Figures S9, S10 and S11). Together, our results show that the approach works best when the training set contains all cell types present in the test set, and when transcriptional differences between cell types are large and stable.

### Cross-validation of newly identified intestinal cell types as an application of *Auto-annot*

Recent studies revealed the intestinal cell type composition utilizing single-cell transcriptomics of intestinal biopsies taken from inflammatory bowel disease (IBD) patients (including ulcerative colitis and Crohn’s disease), healthy donors, or mice, as reviewed recently in [48]. However, the utilized cell type nomenclatures are inconsistent between these studies and various novel cell types were discovered. Here we show how to use *Besca*’s supervised machine learning method *Auto-annot* to cross-validate these disparate cell type annotations. We focus on two major studies: Smillie2019 (human colon epithelium and lamina propria during ulcerative colitis) [33] and Martin2019 (human ileum lamina propria during Crohn’s disease) [34]. In addition, we perform cell type annotation across species using the Haber2017 dataset (mouse small intestine epithelium) [35].

Firstly, we use the Smillie2019 and Martin2019 datasets to train a model with one dataset and apply it to the other, respectively. Both datasets were processed with *Besca*’s standard workflow and cell type annotations were adopted from the respective publications. Both studies provide a coarse cell type annotation (Fig. 5a left) as well as a fine grained cell type annotation (see Supplementary Figure S12 and for Smillie2019 fine-grained fibroblasts Fig. 5d left). Epithelial cell annotations are missing from the Martin2019 author’s annotation, because those cells were excluded in the original study. Therefore, they are labelled as “unknown” in our comparison. The *Auto-annot* module identifies the corresponding cell types in the unseen dataset, respectively (Fig. 5a,b,c and Supplementary Figure S12). Still, there remains some ambiguity mainly within lymphocytes.

**Fig. 5.**
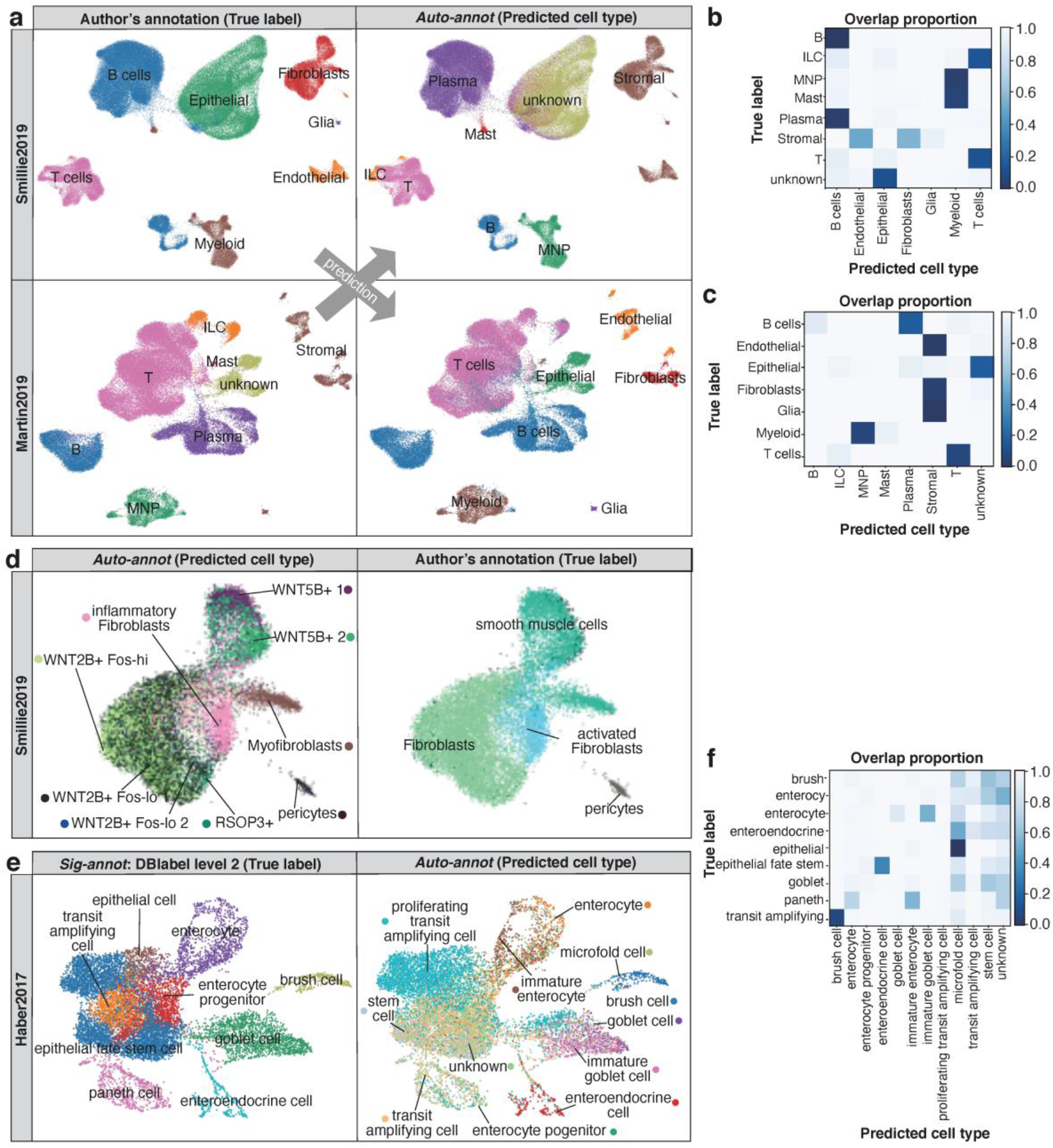
Supervised machine learning to compare intestinal cell type annotations in scRNA-seq data. **a** UMAP representations of the coarse-grained cell types annotated in the Smillie2019 and Martin2019 datasets based on author’s annotations (left) and predictions based on *Besca*’s *Auto-annot* module (right). **b,c** Confusion matrices comparing the true labels from the author’s annotation and the cell types predicted in the Smillie2019 dataset from the Martin2019 annotation (b) and in the Martin2019 dataset predicted from Smillie2019 (c). **d** Discovery of inflammatory or activated fibroblasts point to the same cell community in both studies as exemplified in the Smillie2019 dataset by the author’s annotation (left) and prediction from Martin2019 (right). **e** UMAP representations of the mouse small intestinal epithelial cells from Haber2017 showing the reference DBlabel cell type annotation (left) and cell types predicted from Smillie2019 human colon (right), and **f** the corresponding confusion matrix.

In both studies a new type of disease-relevant fibroblasts was discovered and named inflammatory fibroblasts in ulcerative colitis [33] or activated fibroblasts in Crohn’s disease [34]. Here, we show how our machine learning approach could clearly confirm that these two fibroblast communities belong to the same cell type (Fig. 5d and Supplementary Figure S12). The comparison revealed further differences in the cell type annotation for the enteric nervous systems and B cells, which could be driven by biological differences, experimental differences, or simply different cell type nomenclatures used (Supplementary Figure S12). The results show that our approach can be used to match cell type identities across studies and obtain a more cohesive picture of a tissue’s cell type composition.

Finally, we performed a cross-species comparison. The Haber2017 small intestine mouse dataset includes only epithelial cells and was used as a test dataset. As the training dataset we chose the Smillie2019 human colon dataset and trained the machine learning model on the epithelial cells only. This approach clearly identified enterocytes, enteroendocrine cells, goblet cells and brush (tuft) cells (Fig. 5e,f). The overall gradient from stem and transit amplifying cells to precursor and fully differentiated cells was mainly reproduced, but with less accuracy than the aforementioned discrete cell types (Fig. 5e,f). Paneth cells led to confusion in this scenario, because they are highly abundant in the mouse small intestine [35], but mainly absent in colon and not annotated in the human colon training data [33]. Similar results were achieved by using the fine-grained annotation from Smillie2019 and by the reverse prediction from mouse to human (see Supplementary Figure S13). The results show that a cross-species prediction is generally possible and can provide important insights for translational research.

### scRNA-seq-informed cell deconvolution through *Bescape*

Cell deconvolution aims to estimate cell type proportions from bulk sample transcriptomic data based on cell type specific gene expression profiles (GEPs). Derivation of GEPs relevant for different bulk RNA-seq experiments has remained a challenge. As scRNA-seq data is being collected and annotated at an unprecedented rate, this offers the potential to leverage on the newly gathered knowledge [49]. *Besca*’s deconvolution framework *Bescape* facilitates the usage of established deconvolution methods directly on any scRNA-seq data of choice (Fig. 1e).

Most available tools do not offer the flexibility to introduce user defined cell specific GEPs, instead relying solely on the authors’ carefully curated ones. The application and performance of the cell deconvolution results are then limited to the scope of the tissue and cell types embedded in the curated set. For example, GEPs derived from microarray data from haematological malignancies will have a limited scope of application in deconvoluting cell proportions from bulk RNA-seq sequenced from solid tumour biopsies. In other words, the performance of the results cannot be disentangled between the algorithm itself and the cell type specific gene expression embedded in different tools.

The *Bescape* module aims to leverage on the data collected from the ongoing effort in the understanding of scRNA-seq signals as basis vectors to estimate the cell composition in heterogeneous bulk RNA-seq readouts. As the deconvolution algorithms have made significant progress over the past years [50,51], the focus is now being placed on the specificity of the GEPs that are used as basis vectors to estimate the cell composition addressing platform, tissue and indication variability [52]. This is where *Besca*’s standard workflow and automated cell type annotations from scRNA-seq have a direct impact.

In order to allow for simple incorporation of reference scRNA-seq datasets to generate GEPs for cell types of interest and addressing challenges such as collinearity of closely related cell types, *Bescape* includes two recent cell deconvolution tools, *SCDC* [16] and *MuSiC* [17] (see Methods). Furthermore, as most deconvolution methods are implemented in *R* (https://www.r-project.org/) packages, several steps are needed to run the deconvolution module seamlessly in the background. In short, *Bescape* provides a containerized environment to run the different tools (see also Supplementary Figure S14). More specifically, it first provides a notebook with the combination of *Python* and *R* scripts for the conversion of an *AnnData* h5ad file (https://anndata.readthedocs.io) to an eSet object (https://www.rdocumentation.org/packages/Biobase/topics/eSet) needed to run the deconvolution algorithm in *R*. In addition, *Bescape* provides a notebook in *Python* to run the deconvolution in a *Docker* image (https://docs.docker.com/) based on a user specified reference *eSet* scRNA-seq and a bulk RNAseq dataset.

To extract the information from a reference scRNA-seq dataset, two sets of GEPs are generated from the *Besca* workflow immediately following the cell type annotation step: (1) GEPs can either be generated using all genes from the scRNA-seq reference dataset without performing any feature selection, these are extracted from the functionality provided by *SCDC* and *MuSiC* or (2) based on a subset of highly variable gene expressions defined in the standard workflow. The first set of GEPs is extracted and suitable for use by *MuSiC* and *SCDC* where subsequent weighing of the different genes is performed. The second set of GEPs can be used as input basis matrix for a multitude of cell deconvolution tools such as *EPIC* [53] and *CIBERSORT* [51]. The resulting GEPs derived from the Segerstolpe2016 and Kotliarov2020 datasets (Table 1, [32,37]) utilizing both strategies are shown for comparison in Supplementary Figures S15-S18.

Here, we focus on the first strategy, applied to both datasets, utilizing *SCDC* by example, as it shows how to leverage scRNA-seq data to demonstrate *Bescape*’s functionality best. Bulk RNA-seq was simulated from the pancreatic islets [37] and hematopoietic CITE-seq [32] datasets using the GEPs across all genes from the raw count. The use of simulated bulk RNA-seq, where the ground truth of the in-silico ad-mixture is known, allows validation of the estimated cell proportions (see Methods). The estimated proportions from these simulated data using *SCDC* correlate highly with the ground truth across samples for both datasets (Fig. 6a,b).The estimated proportions show high Pearson correlation with the ground truth, and corresponding low root mean square deviation (RMSD) and mean absolute deviation (mAD), in both tissues for all the cell types that were annotated from the *Besca* workflow (Tables 2 and 3). There are a few exceptions where the cell type GEPs are less well defined (see Supplementary Figures S15 and S17).

**Table 2.** *SCDC* deconvolution results based on simulated bulk RNA-seq from *SCDC* GEPs on pancreatic islets reference scRNA-seq from Segerstolpe2016.

| Cell type | Pearson correlation | RMSD | mAD |
| --- | --- | --- | --- |
| blood vessel endothelial cell | 0.68 | 0.017 | 0.010 |
| enteroendocrine cell | 0.52 | 0.064 | 0.047 |
| fibroblast | 0.87 | 0.020 | 0.017 |
| macrophage | 0.98 | 0.004 | 0.002 |
| pancreatic A cell | 0.97 | 0.043 | 0.033 |
| pancreatic acinar cell | 0.99 | 0.049 | 0.024 |
| pancreatic D cell | 0.88 | 0.023 | 0.018 |
| pancreatic ductal cell | 0.83 | 0.075 | 0.043 |
| PP cell | 0.87 | 0.041 | 0.033 |
| type B pancreatic cell | 0.94 | 0.055 | 0.040 |

**Table 3.** *SCDC* deconvolution results based on simulated bulk RNA-seq from *SCDC* GEPs on hematopoietic reference CITE-seq data from Kotliarov2020.

| Cell type | Pearson correlation | RMSD | mAD |
| --- | --- | --- | --- |
| B cells | 0.98 | 0.009 | 0.007 |
| CD4 T cells | 0.79 | 0.075 | 0.062 |
| CD8 T cells | 0.90 | 0.021 | 0.016 |
| classical monocytes | 0.99 | 0.014 | 0.011 |
| DN T cells | 0.13 | 0.078 | 0.058 |
| ILCs | 0.81 | 0.031 | 0.024 |
| mDCs | 0.75 | 0.009 | 0.008 |
| non-classical monocytes | 0.81 | 0.009 | 0.007 |
| not determined | 0.56 | 0.033 | 0.026 |
| pDCs | 0.61 | 0.003 | 0.002 |

**Fig. 6.**
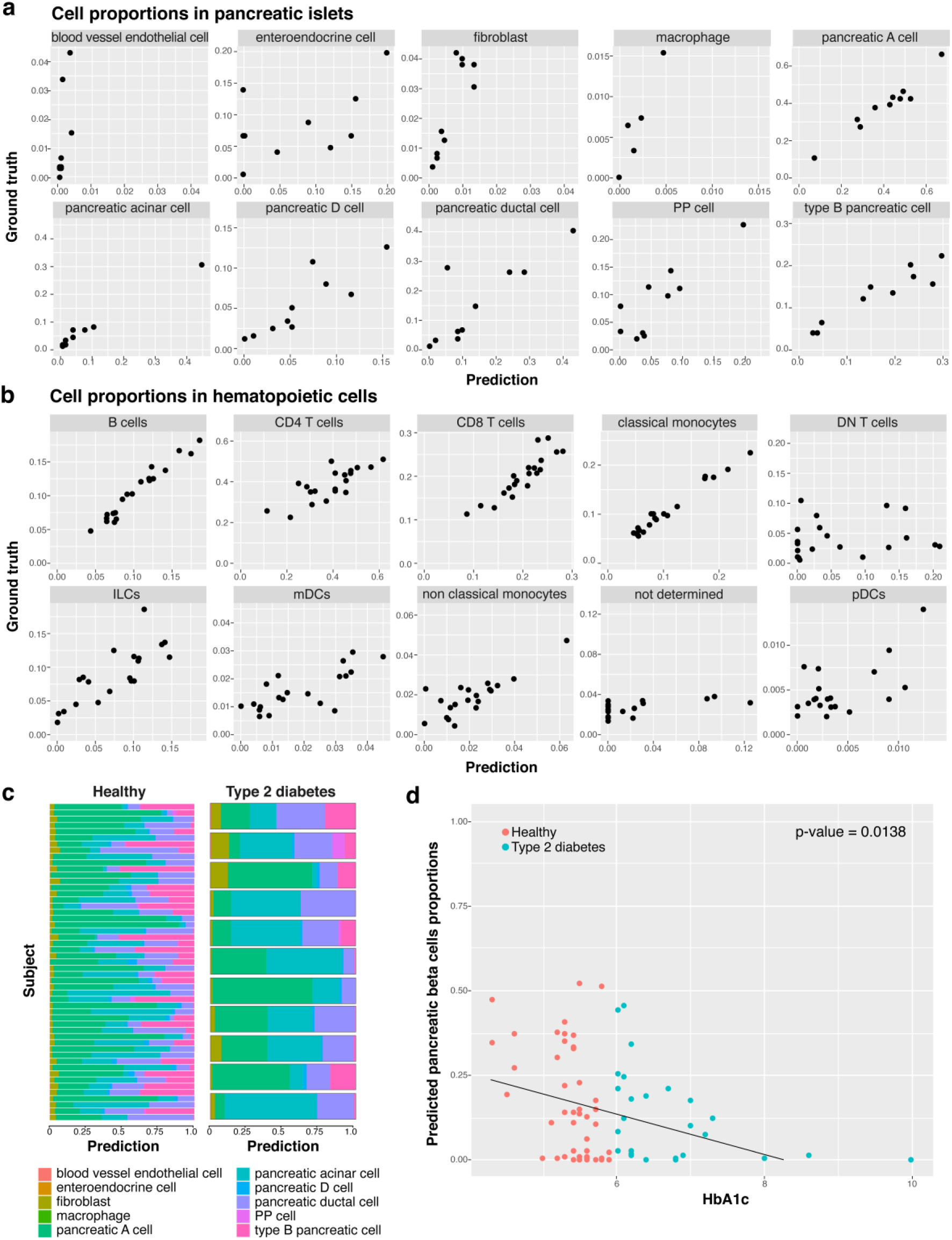
Cell deconvolution using *Bescape*. **a** Measured versus predicted cell proportions in pancreatic islets bulk RNA-seq simulated from Segerstolpe2016. **b** Measured versus predicted cell proportions in hematopoietic bulk RNA-seq simulated from Kotliarov2020. **c** Estimated cell proportions for real pancreatic islets bulk RNA-seq between type 2 diabetes patients and healthy controls. **d** Estimated pancreatic beta cells proportions for real pancreatic islets bulk RNA-seq between type 2 diabetes patients and healthy controls.

Following the *SCDC* manuscript [16], we utilized the study of type 2 diabetes [54] for which the difference in the estimated beta pancreatic cell proportions between type 2 diabetic patients and healthy controls provides a measure for validation of the deconvolution results. Estimated proportions obtained from the real bulk RNA-seq for all 10 cell types using *Besca* cell annotation is shown in Fig. 6c. The estimated pancreatic beta cells are tested and do show the expected lower cell proportions in the type 2 diabetes patients as compared to healthy subjects as shown in Fig. 6d.

It is important to note that the success of cell deconvolution can be measured based on two merits. First, on the accuracy to known proportions estimated based on a known or proxy ground truth either on simulated bulk RNA-seq data or from matched samples measured with more traditional single-cell means (e.g. immunohistochemistry or flow cytometry). Although this is the preferred measure of success, validating the results compared to a ground truth obtained using known cell types can be difficult as these more traditional methods for studying cell heterogeneity rely on a limited repertoire of markers of known cell types. Secondly, the success can also be measured based on the results obtained from embedding estimated cell proportions as covariates in prognostic and predictive models. Indication of success here is harder to determine a priori and would need to be carefully investigated to avoid overfitting.

## Discussion

No two cells are identical; neither are two scRNA-seq experiments. Cells are extracted from different tissues, treated according to lab-specific protocols, and sequenced with a variety of technologies. Still, the vast amount of available scRNA-seq studies provokes the ambition to reuse the valuable experimental data and to re-assess them by comparing between studies. Streamlined and standardized workflows, such as those presented here, strive to find balance between automation and flexibility, which brings efficiency, reusability, and reproducibility of data processing. The loss in flexibility is compensated by bringing scRNA-seq results to a level that allows for cross-study comparisons and finally integration into larger cell atlases.

*Besca*’s *Sig-annot* module implements such a process by automatizing cell type annotation through an unsupervised clustering approach followed by signature-based hierarchical cell type identification. It mirrors the manual annotation approach, but enforces a harmonized annotation schema and hence guarantees comparability between studies. It also captures knowledge of cell type markers that is gained in this process in explicit gene signatures that can be easily shared, re-assessed and improved across different conditions, studies, and technologies. This signature-based approach is valuable for specific tissues and disease phenotypes as an approach to harmonize annotations across various cell atlases, which is critical for holistic disease understanding (see e.g. [1,48,55]). Still, each tissue and fine-grained cell type needs to be incorporated and optimized for in the annotation schema.

In contrast, supervised approaches overcome certain challenges faced by unsupervised clustering [56] and therefore generalize better. Importantly, they allow for the utilization of curated high-quality annotations by transferring them to new studies efficiently. Such approaches not only allow for the comparison of cell annotations between studies, but even across species. They depend on well annotated reference datasets containing harmonized cell type annotations. We expect cell atlas projects (see e.g. [55,57–61]) to provide such annotations in the near future for all major tissues, which would allow for a wide applicability of supervised approaches. Furthermore, the cell type annotation can also be resolved with correlation-based approaches such as *singleR* [62], *scMCA* [63], or *SCMAP* [64].

Various examples in this manuscript and previous studies show that the automation of cell type annotation is feasible to a certain extent and technically not too complex. As cells can be grouped by multiple orthogonal criteria such as surface markers, functions, cell cycle states, differentiation stages, or activation levels, a clear definition of concrete cell types remains controversial and a more general concept of cell types will be needed in the future [29]. Setting aside the controversy in cell type definition, our work already provides tools and best practices to achieve better reference cell annotations and to share the gene signatures that capture the knowledge about how they were derived. Like the human reference genome (which does not ultimately reflect a human genome consensus [65] and still serves many practical purposes) accelerated genomic research, such reference cell type annotations will accelerate our understanding of biological systems even though they reflect only a subset of a cell’s characteristics.

Finally, these cell type definitions help investigate changes in cell composition and differentially expressed genes within certain cell types, which are often postulated as indications of disease progression or response to stimulation and perturbation [66]. While scRNA-seq offers the possibility to investigate these hypotheses, the current cost as well as the technical and logistical challenges associated with the technology are preventing large scale studies [67], particularly in a clinical trial setting. Although this is likely to improve over time as the technology matures, large numbers of biological replicates are currently measured using bulk RNA-seq. In these samples, heterogeneity resulting from the distinct cell type composition of the probed material can often confound the signals, making it difficult to interpret results. By leveraging annotated reference scRNAseq datasets in combination with cell deconvolution methods, the cell composition of bulk RNA-seq samples can be robustly estimated. This information can then either be used directly as biomarkers or as covariates towards inferring more robust differential gene expression results.

## Conclusions

In sum, the core benefits of adopting *Besca* for scRNA-seq data analysis are automation, standardization, and reusability. This is achieved (1) by a generalized standard workflow, including CITE-seq data processing, (2) by the automation of the cell type annotation process with two complementary approaches, (3) by managing knowledge about cell type marker gene signatures in *GeMS*, (4) by informing deconvolution algorithms to make better use of bulk transcriptional data, and (5) by building upon the widely used *Scanpy* toolkit. We expect that *Besca*, published as an open-source software contribution to the community, will promote interoperability, reusability, and interpretability of scRNA-seq data. Finally, *Besca* will be part of the many components that pave the way for a reference catalogue of cell types and their reactions to various perturbations. This catalogue will allow a deeper understanding of human diseases and their interventions.

## Methods

### Example data

The following publicly available single-cell datasets from ten studies were reprocessed (see also Table 1). Three datasets cover blood- and bone-marrow-derived hematopoietic cells:

- PBMC3k (https://doi.org/10.5281/zenodo.3948150) includes healthy peripheral blood mononuclear cell (PBMC) samples from one donor, a reference dataset often used in single-cell tutorials based on 10X Genomics data (https://www.10xgenomics.com/).
- Granja2019 (https://doi.org/10.5281/zenodo.3944753) includes bone marrow mononuclear cell (BMMCs) and PBMC samples from healthy donors [31]. In addition to scRNA-seq, several protein markers were also probed by CITE-seq.
- Kotliarov2020 (https://doi.org/10.5281/zenodo.3938290) includes baseline PBMC samples from healthy donors, who were high and low responders to influenza vaccines [32]. In addition to scRNA-seq, a high number of protein markers were also probed by CITE-seq.

Four datasets reveal the intestinal cell composition:

- Smillie2019 (https://doi.org/10.5281/zenodo.3960617) includes colon epithelium and lamina propria samples from healthy donors and ulcerative colitis patients [33].
- Martin2019 (https://doi.org/10.5281/zenodo.3862132) includes ileal lamina propria samples from Crohn’s disease patients [34].
- Haber2017 (https://doi.org/10.5281/zenodo.3935782) includes murine small intestine samples [35].
- Lee2020 (https://doi.org/10.5281/zenodo.3967538) includes tumor and non-malignant colon samples from colorectal cancer (CRC) patients [36].

Three datasets are pancreas-derived:

- Segerstolpe2016 (https://doi.org/10.5281/zenodo.3928276) includes pancreatic islet cells from healthy donors and type 2 diabetic patients [37].
- Peng2019 (https://doi.org/10.5281/zenodo.3969339) includes tumor and non-malignant pancreatic samples from pancreatic ductal adenocarcinoma (PDAC) and non-pancreatic tumor patients [38].
- Baron2016 (https://doi.org/10.5281/zenodo.3968315) includes pancreatic samples from healthy donors [39].

### *Besca*’s standard workflow

*Besca*’s standard workflow starts with loading the count matrix obtained from a preprocessing pipeline (demultiplexing, read alignment, feature counting), and the annotation of the matrix, including barcodes, genes and, if available metadata associated to the datasets, including biological (e.g. donor, experimental condition) and technical (e.g. batches, protocols differences) variables. Before proceeding with analysis, quality control (QC) is performed. This includes visualizing drop-outs and sequencing saturation as well as performing cell and gene filtering. During cell filtering all barcodes that do not correspond to viable cells are removed. Cell filtering is performed on the basis of three QC covariates: the number of counts per barcode, the number of genes per barcode, and the contribution from mitochondrial genes per barcode. Each of the covariates are examined for outliers by thresholding as described in [11]. During gene filtering, transcripts which are only expressed in a few cells are removed to reduce dataset dimensionality. As recommended by Luecken and Theis [11], the filtering threshold for genes should be set to the minimum cell cluster size that is of interest. As QC filtering is highly dependent on the dataset the filtering thresholds need to be defined by the user before running the workflow. Correctly chosen thresholds are verified through knee-plot graphics within the pipeline.

After QC, the expression values are normalized. Normalization is performed using count depth scaling and count values are log(x+1)-transformed. To reduce dataset dimensionality before clustering, the highly variable genes within the dataset are selected. By default, genes are defined as being highly variable when they have a minimum mean expression of 0.0125, a maximum mean expression of 3 and a minimum dispersion of 0.5. Technical variance is removed by regressing out the effects of count depth and mitochondrial gene content and the gene expression values are scaled to a mean of 0 and variance of 1 with a maximum value of It needs to be mentioned here that correction of mitochondrial gene content might not be considered a technical variance correction but removal of biological variability. If this correction is not desired, the threshold for mitochondrial gene content correction can be set to 1. Based on the best practices suggested by Luecken and Theis, technical variance should be corrected before selection of highly variable genes. In *Besca*’s standard workflow though this order is reversed, due to regress-out being a very time-consuming computational process which can be significantly sped up by only calculating corrected values for the previously selected highly variable genes. For larger datasets it is absolutely essential to reduce dimensionality beforehand for regress-out to even complete.

Finally, dimensionality reduction and clustering is performed. The first 50 principle components are calculated and used as input for calculation of the 10 nearest neighbours. The neighbourhood graph is then embedded into two-dimensional space using the UMAP (Uniform Manifold Approximation and Projection) algorithm [68]. Cell communities are detected using the Leiden algorithm [69] at a resolution of 1 by default.

For CITE-seq data, the protein marker abundance values are loaded separately to the gene expression values and stored in its own data object. Previously determined cell barcode filtering to identify viable cells on the basis of gene expression values is applied to the CITE-seq data. Unlike gene expression counts, protein marker counts are normalized using centred log ratios. If less than 50 markers were measured the entire count matrix is used as input for the nearest neighbour calculation otherwise, as in the gene expression data, the first 50 principal components are calculated. The rest of the CITE-seq pipeline is analogous to the gene expression pipeline. At the end of the workflow the results are homogenized into one data object which contains clustering and visualization results of both gene expression and protein abundance from CITE-seq data.

Analysis results are exported into interoperable file formats to allow FAIR data management of analysis results. This includes the Matrix Market exchange format (https://math.nist.gov/MatrixMarket/formats.html) for sparse count matrices, GCT (https://software.broadinstitute.org/software/igv/GCT) for dense count matrices, and simple tab-separated or comma-separated values formats for metadata and as interface for the cell deconvolution package *Bescape*, respectively. Clustering results or cell type labelling can be exported including pre-computed average expression and ranked marker gene lists per cluster or cell type.

### Annotation of cell types based on CITE-seq data

A fine-grained annotation of the cells contained within the Kotliarov2020 dataset [32] was generated on the basis of the labelled protein antibody counts from CITE-seq. The normalized protein counts were exported to FCS files using the R package flowCore [70,71] (R package version 2.0.1) and loaded into FlowJo™ Software (FlowJo™ Software Mac Version 10.6.2.

Ashland, OR: Becton, Dickinson and Company; 2019). The gating strategy used to identify individual cell populations is outlined in Supplementary Figure S2. Gating of individual cell populations was based on the gating strategy utilized in [72]. Barcodes from identified cell populations were exported from FlowJo™ Software to csv files and loaded into *Besca* for visualization.

### Sig-annot, signature-based automated cell type annotation

The annotation process has three components:

1. a nomenclature table with long and short names, according to *Cell Ontology* [40]

- see Supplementary Table S1 and https://github.com/bedapub/besca/blob/master/besca/datasets/nomenclature/CellTypes_v1.tsv
2. a configuration file including all the cell types to be considered, their parent (or "none"), a factor to be multiplied with the cut-off for scoring a cluster positive or negative for the signature based on the Mann-Whitney test and the order in which to consider the signatures (only first positive one matching a cluster will be taken into account). Two distinct default configuration files are provided with *Besca*, covering mouse and humans. Users are free to adjust the parameters in the files, and tailor these according to tissues or dataset.

- Human: Supplementary Table S2 and https://github.com/bedapub/besca/blob/master/besca/datasets/genesets/CellNames_scseqCMs6_config.tsv
- Mouse: Supplementary Table S3 and https://github.com/bedapub/besca/blob/master/besca/datasets/genesets/CellNames_scseqCMs6_config.mouse.tsv
3. a GMT file with the signatures, in line with the nomenclature table.

- see Supplementary Table S4 and https://github.com/bedapub/besca/blob/master/besca/datasets/genesets/CellNames_scseqCMs6_sigs.gmt

### *Auto-annot*, supervised automated cell type annotation

Besca’s *Auto-annot* module, a supervised machine learning workflow, can be run independently from the standard workflow and works as follows:

- Initially the training datasets are merged to form a combined training dataset using Scanorama [73], in the case where multiple training datasets are available, and complemented with the testing dataset. A parameter specifies if the resulting integrated gene expression matrix contains the intersection of all genes, the intersection of previously selected highly variable genes, or genes of a previously defined signature.
- Secondly, the *Python* package *scikit-learn* (https://scikit-learn.org) is used to train a classifier based on the merged training datasets. Two classification approaches are implemented, SVM and logistic regression. For SVM, one can choose between SVM with linear kernel (*linear*); SVM with linear kernel using stochastic gradient descent (*sgd*); SVM with radial basis function kernel (*rbf*), which should be used on small datasets only due to longer runtime. For logistic regression, the options are multinomial loss (*logistic_regression*); logistic regression with one versus rest classification, without normalised probability scores (*logistic_regression_ovr*); logistic regression with elastic loss, cross validated among multiple l1 ratio (*logistic_regression_elastic*). We recommend *logistic_regression* as default option.
- Finally, the fitted model is used to predict cell types in the test dataset and predictions are added to the metadata. A probability threshold can be defined for logistic regression classifiers, to classify only cells reaching the defined threshold. In order to compare the predicted cell types to a ground truth already annotated in the test datasets, a report can be generated including precision, recall, and F1 metrics as well as confusion matrix and automatically annotated UMAP plots.

### *Bescape,* cell deconvolution

At the core of the cell deconvolution algorithm is a regression based problem. The concept is not novel, it has already been investigated for microarray data [74]. The combination of how newly derived cell specific GEP from scRNA-seq data can be used is the key factor that has evolved considerably over time. At a broad level, there are two categories of cell deconvolution, it is either a *full deconvolution* where neither the source nor the mixing process is known or a *partial deconvolution* where there is priori knowledge of the sources or the mixing process. Although a completely unsupervised approach can be taken, where the non-negative matrix factorization is suitable, it has been proven to show low accuracy and difficulty in handling the collinearity of the genes [16]. The research focus is placed on partial deconvolution with known signatures used as bases to estimate the proportions in the bulk tissue. Such approaches have been developed using constrained least squares regression (*EPIC*) [53] and *v*-support vector regression (*CIBERSORT*) [51]. These methods either use microarray or a mixture of bulk RNA and scRNA-seq data to build a single GEP as a basis vector. Two distinct sets of cell type specific GEPs are generated as part of the toolkit. Equipped with the derived GEPs, the users will have the choice to apply the deconvolution algorithm of their choice.

Two recent methods have been included in the cell deconvolution module allowing for direct incorporation of reference scRNA-seq datasets and addressing some of the shortcomings of previous methods. The first scRNA-seq reference dataset based method is *MuSiC* [17]. In short, this method uses a constrained least square regression but factors in the weighing of the different genes to reduce the impact of the residuals on the fit from genes that are less informative in terms of cell types differentiation and thus, eliminates the need for preselection of genes. Most importantly, it addresses the hierarchical nature of cell lineages with a recursive tree guided search, similar to gating strategy in FACS, by first grouping similar cell types into the same cluster and estimating cluster proportions, then recursively repeating the previous step within each cluster identified. At each recursion stage, the focus is only on differentially expressed genes across cell types within the cluster. Consequently, the residuals are determined on only the subset of genes important to differentiate the cells within the cluster as opposed to being diluted by genes which share a common profile.

The second method included in the *Bescape* module is *SCDC* [16], an ensemble approach allowing for multiple scRNA-seq reference datasets. In short, similar to *MuSiC*, a weighted non-negative least square regression is adopted but differs slightly on how the weights are assigned to the genes. The salient point of the method is an additional layer of abstraction being introduced by assigning different weights for each reference scRNA-seq dataset. Higher weights are attributed to reference datasets that can fit the gene expression profiles of bulk RNA-seq samples better based on defined performance metric.

### Generating simulated bulk

Simulated bulk RNA-seq was generated to evaluate the estimated proportions of the selected cell types with ground truth from a known in-silico mixture. The annotated scRNA-seq data can be used directly by *SCDC* and *MuSiC* where no user specified feature selection based on marker genes is needed, instead a higher weight is assigned to features showing high variability across annotated cell types and low variability across samples [16,17]. The simulated bulk is based on linear regressions where the cell fractions (weights) are taken from a uniform distribution, thus without factoring in any prior knowledge of the range of cell proportions of the different cell types, and scaled for the total to add up to 1. The GEPs of the cell types constitute the basis matrix needed to construct the bulk RNA-seq vector. This step is repeated for several instances representing different subjects’ bulk RNA-seq data.

## Notes

This publication is part of the Human Cell Atlas: www.humancellatlas.org/publications.

## References

1. Muus C, Luecken MD, Eraslan G, Waghray A, Heimberg G, Sikkema L, et al. Integrated analyses of single-cell atlases reveal age, gender, and smoking status associations with cell type-specific expression of mediators of SARS-CoV-2 viral entry and highlights inflammatory programs in putative target cells. bioRxiv. Cold Spring Harbor Laboratory; 2020;2020.04.19.049254.

2. Ziegler CGK, Allon SJ, Nyquist SK, Mbano IM, Miao VN, Tzouanas CN, et al. SARS-CoV-2 Receptor ACE2 Is an Interferon-Stimulated Gene in Human Airway Epithelial Cells and Is Detected in Specific Cell Subsets across Tissues. Cell. 2020;181:1016-1035.e19.

3. Kim J, Koo B-K, Knoblich JA. Human organoids: model systems for human biology and medicine. Nature Reviews Molecular Cell Biology. Nature Publishing Group; 2020;1–14.

4. Zhang Q, Caudle WM, Pi J, Bhattacharya S, Andersen ME, Kaminski NE, et al. Embracing Systems Toxicology at Single-Cell Resolution. Curr Opin Toxicol. 2019;16:49–57.

5. Efremova M, Vento-Tormo M, Teichmann SA, Vento-Tormo R. CellPhoneDB: inferring cell–cell communication from combined expression of multi-subunit ligand–receptor complexes. Nature Protocols. Nature Publishing Group; 2020;15:1484–506.

6. Szabo PA, Levitin HM, Miron M, Snyder ME, Senda T, Yuan J, et al. Single-cell transcriptomics of human T cells reveals tissue and activation signatures in health and disease. Nature Communications. Nature Publishing Group; 2019;10:4706.

7. Saelens W, Cannoodt R, Todorov H, Saeys Y. A comparison of single-cell trajectory inference methods. Nature Biotechnology. Nature Publishing Group; 2019;37:547–54.

8. Lee JTH, Hemberg M. Supervised clustering for single-cell analysis. Nature Methods. 2019;16:965–6.

9. Miao Z, Moreno P, Huang N, Papatheodorou I, Brazma A, Teichmann SA. Putative cell type discovery from single-cell gene expression data. Nature Methods. Nature Publishing Group; 2020;17:621–8.

10. Hie B, Peters J, Nyquist SK, Shalek AK, Berger B, Bryson BD. Computational Methods for Single-Cell RNA Sequencing. Annual Review of Biomedical Data Science. 2020;3:339–64.

11. Luecken MD, Theis FJ. Current best practices in single-cell RNA-seq analysis: a tutorial. Molecular Systems Biology. John Wiley & Sons, Ltd; 2019;15:e8746.

12. Stegle O, Teichmann SA, Marioni JC. Computational and analytical challenges in single-cell transcriptomics. Nat Rev Genet. 2015;16:133–45.

13. Angerer P, Simon L, Tritschler S, Wolf FA, Fischer D, Theis FJ. Single cells make big data: New challenges and opportunities in transcriptomics. Current Opinion in Systems Biology. 2017;4:85–91.

14. Svensson V, Vento-Tormo R, Teichmann SA. Exponential scaling of single-cell RNA-seq in the past decade. Nature Protocols. Nature Publishing Group; 2018;13:599–604.

15. Wolf FA, Angerer P, Theis FJ. SCANPY: large-scale single-cell gene expression data analysis. Genome Biology. 2018;19:15.

16. Dong M, Thennavan A, Urrutia E, Li Y, Perou CM, Zou F, et al. SCDC: bulk gene expression deconvolution by multiple single-cell RNA sequencing references. Brief Bioinformatics. 2020;

17. Wang X, Park J, Susztak K, Zhang NR, Li M. Bulk tissue cell type deconvolution with multi-subject single-cell expression reference. Nature Communications. Nature Publishing Group; 2019;10:380.

18. Stoeckius M, Hafemeister C, Stephenson W, Houck-Loomis B, Chattopadhyay PK, Swerdlow H, et al. Simultaneous epitope and transcriptome measurement in single cells. Nature Methods. Nature Publishing Group; 2017;14:865–8.

19. Bergen V, Lange M, Peidli S, Wolf FA, Theis FJ. Generalizing RNA velocity to transient cell states through dynamical modeling. Nature Biotechnology. Nature Publishing Group; 2020;1–7.

20. Setty M, Kiseliovas V, Levine J, Gayoso A, Mazutis L, Pe’er D. Characterization of cell fate probabilities in single-cell data with Palantir. Nat Biotechnol. 2019;37:451–60.

21. Sturm G, Szabo T, Fotakis G, Haider M, Rieder D, Trajanoski Z, et al. Scirpy: A Scanpy extension for analyzing single-cell T-cell receptor sequencing data. Bioinformatics. 2020;

22. Avila Cobos F, Vandesompele J, Mestdagh P, De Preter K. Computational deconvolution of transcriptomics data from mixed cell populations. Bioinformatics. 2018;34:1969–79.

23. Wilkinson MD, Dumontier M, Aalbersberg IjJ, Appleton G, Axton M, Baak A, et al. The FAIR Guiding Principles for scientific data management and stewardship. Scientific Data. Nature Publishing Group; 2016;3:160018.

24. Jassal B, Matthews L, Viteri G, Gong C, Lorente P, Fabregat A, et al. The reactome pathway knowledgebase. Nucleic Acids Res. 2020;48:D498–503.

25. Wang Z, Monteiro CD, Jagodnik KM, Fernandez NF, Gundersen GW, Rouillard AD, et al. Extraction and analysis of signatures from the Gene Expression Omnibus by the crowd. Nature Communications. Nature Publishing Group; 2016;7:12846.

26. Zhang X, Lan Y, Xu J, Quan F, Zhao E, Deng C, et al. CellMarker: a manually curated resource of cell markers in human and mouse. Nucleic Acids Res. Oxford Academic; 2019;47:D721–8.

27. Subramanian A, Tamayo P, Mootha VK, Mukherjee S, Ebert BL, Gillette MA, et al. Gene set enrichment analysis: a knowledge-based approach for interpreting genome-wide expression profiles. Proc Natl Acad Sci USA. 2005;102:15545–50.

28. Liberzon A, Birger C, Thorvaldsdóttir H, Ghandi M, Mesirov JP, Tamayo P. The Molecular Signatures Database Hallmark Gene Set Collection. Cell Systems. 2015;1:417–25.

29. Wagner A, Regev A, Yosef N. Revealing the vectors of cellular identity with single-cell genomics. Nature Biotechnology. 2016;34:1145–60.

30. Abdelaal T, Michielsen L, Cats D, Hoogduin D, Mei H, Reinders MJT, et al. A comparison of automatic cell identification methods for single-cell RNA sequencing data. Genome Biology. 2019;20:194.

31. Granja JM, Klemm S, McGinnis LM, Kathiria AS, Mezger A, Corces MR, et al. Single-cell multiomic analysis identifies regulatory programs in mixed-phenotype acute leukemia. Nature Biotechnology. Nature Publishing Group; 2019;37:1458–65.

32. Kotliarov Y, Sparks R, Martins AJ, Mulè MP, Lu Y, Goswami M, et al. Broad immune activation underlies shared set point signatures for vaccine responsiveness in healthy individuals and disease activity in patients with lupus. Nature Medicine. Nature Publishing Group; 2020;26:618–29.

33. Smillie CS, Biton M, Ordovas-Montanes J, Sullivan KM, Burgin G, Graham DB, et al. Intra- and Inter-cellular Rewiring of the Human Colon during Ulcerative Colitis. Cell. Elsevier; 2019;178:714–730.e22.

34. Martin JC, Chang C, Boschetti G, Ungaro R, Giri M, Grout JA, et al. Single-Cell Analysis of Crohn’s Disease Lesions Identifies a Pathogenic Cellular Module Associated with Resistance to Anti-TNF Therapy. Cell. Elsevier; 2019;178:1493–1508.e20.

35. Haber AL, Biton M, Rogel N, Herbst RH, Shekhar K, Smillie C, et al. A single-cell survey of the small intestinal epithelium. Nature. Nature Publishing Group; 2017;551:333–9.

36. Lee H-O, Hong Y, Etlioglu HE, Cho YB, Pomella V, Van den Bosch B, et al. Lineage-dependent gene expression programs influence the immune landscape of colorectal cancer. Nature Genetics. Nature Publishing Group; 2020;52:594–603.

37. Segerstolpe Å, Palasantza A, Eliasson P, Andersson E-M, Andréasson A-C, Sun X, et al. Single-Cell Transcriptome Profiling of Human Pancreatic Islets in Health and Type 2 Diabetes. Cell Metabolism. Elsevier; 2016;24:593–607.

38. Peng J, Sun B-F, Chen C-Y, Zhou J-Y, Chen Y-S, Chen H, et al. Single-cell RNA-seq highlights intra-tumoral heterogeneity and malignant progression in pancreatic ductal adenocarcinoma. Cell Research. Nature Publishing Group; 2019;29:725–38.

39. Baron M, Veres A, Wolock SL, Faust AL, Gaujoux R, Vetere A, et al. A Single-Cell Transcriptomic Map of the Human and Mouse Pancreas Reveals Inter- and Intra-cell Population Structure. cels. Elsevier; 2016;3:346–360.e4.

40. Diehl AD, Meehan TF, Bradford YM, Brush MH, Dahdul WM, Dougall DS, et al. The Cell Ontology 2016: enhanced content, modularization, and ontology interoperability. Journal of Biomedical Semantics. 2016;7:44.

41. Malone J, Holloway E, Adamusiak T, Kapushesky M, Zheng J, Kolesnikov N, et al. Modeling sample variables with an Experimental Factor Ontology. Bioinformatics. Oxford Academic; 2010;26:1112–8.

42. Zhang Y, Gao S, Xia J, Liu F. Hematopoietic Hierarchy –An Updated Roadmap. Trends in Cell Biology. Elsevier; 2018;28:976–86.

43. Pliner HA, Shendure J, Trapnell C. Supervised classification enables rapid annotation of cell atlases. Nature Methods. Nature Publishing Group; 2019;16:983–6.

44. Zhang AW, O’Flanagan C, Chavez EA, Lim JLP, Ceglia N, McPherson A, et al. Probabilistic cell-type assignment of single-cell RNA-seq for tumor microenvironment profiling. Nature Methods. Nature Publishing Group; 2019;16:1007–15.

45. Li C, Liu B, Kang B, Liu Z, Liu Y, Chen C, et al. SciBet as a portable and fast single cell type identifier. Nature Communications. Nature Publishing Group; 2020;11:1818.

46. Lin Y, Cao Y, Kim HJ, Salim A, Speed TP, Lin DM, et al. scClassify: sample size estimation and multiscale classification of cells using single and multiple reference. Molecular Systems Biology. John Wiley & Sons, Ltd; 2020;16:e9389.

47. Köhler ND, Büttner M, Theis FJ. Deep learning does not outperform classical machine learning for cell-type annotation. bioRxiv. Cold Spring Harbor Laboratory; 2019;653907.

48. Bigaeva E, Uniken Venema WTC, Weersma RK, Festen EAM. Understanding human gut diseases at single-cell resolution. Hum Mol Genet. 2020;

49. Sturm G, Finotello F, Petitprez F, Zhang JD, Baumbach J, Fridman WH, et al. Comprehensive evaluation of transcriptome-based cell-type quantification methods for immuno-oncology. Bioinformatics. Oxford Academic; 2019;35:i436–45.

50. Gaujoux R, Seoighe C. CellMix: a comprehensive toolbox for gene expression deconvolution. Bioinformatics. Oxford Academic; 2013;29:2211–2.

51. Newman AM, Liu CL, Green MR, Gentles AJ, Feng W, Xu Y, et al. Robust enumeration of cell subsets from tissue expression profiles. Nature Methods. Nature Publishing Group; 2015;12:453–7.

52. Schelker M, Feau S, Du J, Ranu N, Klipp E, MacBeath G, et al. Estimation of immune cell content in tumour tissue using single-cell RNA-seq data. Nature Communications. Nature Publishing Group; 2017;8:2032.

53. Racle J, de Jonge K, Baumgaertner P, Speiser DE, Gfeller D. Simultaneous enumeration of cancer and immune cell types from bulk tumor gene expression data. Valencia A, editor. eLife. eLife Sciences Publications, Ltd; 2017;6:e26476.

54. Fadista J, Vikman P, Laakso EO, Mollet IG, Esguerra JL, Taneera J, et al. Global genomic and transcriptomic analysis of human pancreatic islets reveals novel genes influencing glucose metabolism. PNAS. National Academy of Sciences; 2014;111:13924–9.

55. Ecker JR, Geschwind DH, Kriegstein AR, Ngai J, Osten P, Polioudakis D, et al. The BRAIN Initiative Cell Census Consortium: Lessons Learned toward Generating a Comprehensive Brain Cell Atlas. Neuron. Elsevier; 2017;96:542–57.

56. Kiselev VY, Andrews TS, Hemberg M. Challenges in unsupervised clustering of single-cell RNA-seq data. Nature Reviews Genetics. Nature Publishing Group; 2019;20:273–82.

57. Regev A, Teichmann SA, Lander ES, Amit I, Benoist C, Birney E, et al. The Human Cell Atlas. Gingeras TR, editor. eLife. eLife Sciences Publications, Ltd; 2017;6:e27041.

58. Han X, Zhou Z, Fei L, Sun H, Wang R, Chen Y, et al. Construction of a human cell landscape at single-cell level. Nature. Nature Publishing Group; 2020;581:303–9.

59. Han X, Wang R, Zhou Y, Fei L, Sun H, Lai S, et al. Mapping the Mouse Cell Atlas by Microwell-Seq. Cell. 2018;172:1091–1107.e17.

60. Schaum N, Karkanias J, Neff NF, May AP, Quake SR, Wyss-Coray T, et al. Single-cell transcriptomics of 20 mouse organs creates a Tabula Muris. Nature. Nature Publishing Group; 2018;562:367–72.

61. Snyder MP, Lin S, Posgai A, Atkinson M, Regev A, Rood J, et al. The human body at cellular resolution: the NIH Human Biomolecular Atlas Program. Nature. Nature Publishing Group; 2019;574:187–92.

62. Aran D, Looney AP, Liu L, Wu E, Fong V, Hsu A, et al. Reference-based analysis of lung single-cell sequencing reveals a transitional profibrotic macrophage. Nat Immunol. 2019;20:163–72.

63. Sun H, Zhou Y, Fei L, Chen H, Guo G. scMCA: A Tool to Define Mouse Cell Types Based on Single-Cell Digital Expression. Methods Mol Biol. 2019;1935:91–6.

64. Kiselev VY, Yiu A, Hemberg M. scmap: projection of single-cell RNA-seq data across data sets. Nature Methods. Nature Publishing Group; 2018;15:359–62.

65. Hatje K, Mühlhausen S, Simm D, Kollmar M. The Protein-Coding Human Genome: Annotating High-Hanging Fruits. BioEssays. 2019;41:1900066.

66. Wang Y, Navin NE. Advances and Applications of Single Cell Sequencing Technologies. Mol Cell. 2015;58:598–609.

67. Haghverdi L, Lun ATL, Morgan MD, Marioni JC. Batch effects in single-cell RNA-sequencing data are corrected by matching mutual nearest neighbors. Nat Biotechnol. 2018;36:421–7.

68. McInnes L, Healy J, Melville J. UMAP: Uniform Manifold Approximation and Projection for Dimension Reduction. arXiv:180203426 [cs, stat] [Internet]. 2018 [cited 2020 Aug 2]; Available from: http://arxiv.org/abs/1802.03426

69. Traag VA, Waltman L, van Eck NJ. From Louvain to Leiden: guaranteeing well-connected communities. Scientific Reports. Nature Publishing Group; 2019;9:5233.

70. Hahne F, LeMeur N, Brinkman RR, Ellis B, Haaland P, Sarkar D, et al. flowCore: a Bioconductor package for high throughput flow cytometry. BMC Bioinformatics. 2009;10:106.

71. Ellis B, Haal P, Hahne F, Meur NL, Gopalakrishnan N, Spidlen J, et al. flowCore: flowCore: Basic structures for flow cytometry data [Internet]. Bioconductor version: Release (3.11); 2020 [cited 2020 Aug 2]. Available from: https://bioconductor.org/packages/flowCore/

72. Waugh KA, Araya P, Pandey A, Jordan KR, Smith KP, Granrath RE, et al. Mass Cytometry Reveals Global Immune Remodeling with Multi-lineage Hypersensitivity to Type I Interferon in Down Syndrome. Cell Reports. 2019;29:1893–1908.e4.

73. Hie B, Bryson B, Berger B. Efficient integration of heterogeneous single-cell transcriptomes using Scanorama. Nature Biotechnology. Nature Publishing Group; 2019;37:685–91.

74. Abbas AR, Wolslegel K, Seshasayee D, Modrusan Z, Clark HF. Deconvolution of Blood Microarray Data Identifies Cellular Activation Patterns in Systemic Lupus Erythematosus. PLOS ONE. Public Library of Science; 2009;4:e6098.

