## Supplementary Material and Figures for "*Besca*, a single-cell transcriptomics analysis toolkit to accelerate translational research"

### Standard workflow

#### Supplementary Figure S1

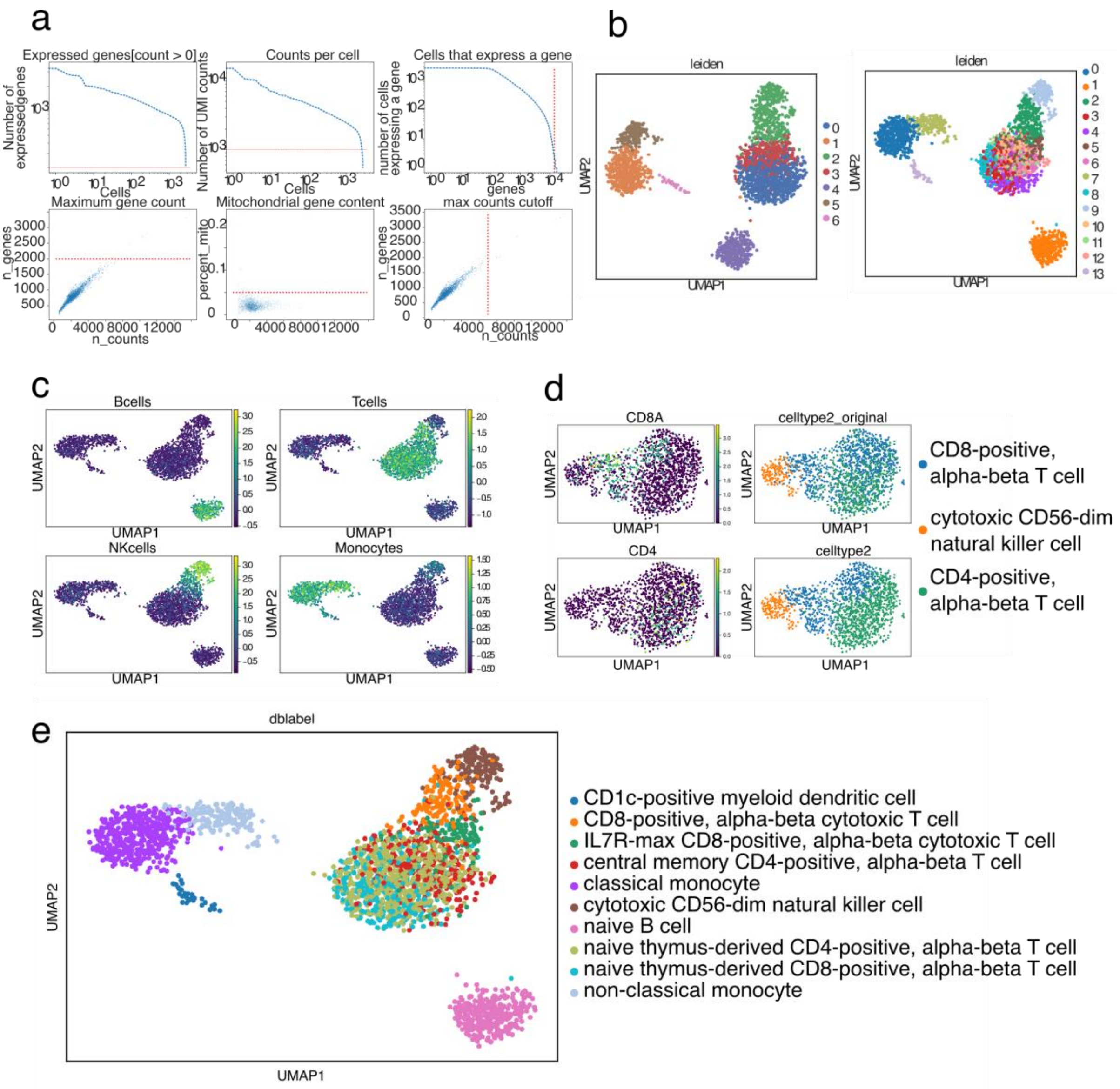

**Supplementary Figure S1** Analysis of the PBMC3k dataset. **a** QC plots and thresholds, the default threshold were lowered. **b** UMAP showing the two clusterings obtained: with default resolution (left) and after increasing the resolution (right). Leiden clustering, default resolution was increased as it was observed that one cluster was mixed in T cells and NK cells (see c). **c** Signature observation. **d** UMAPs of the re-clustered cells: CD8A and CD4 expression (left), top-right UMAP shows cluster assignment after reclustering, bottom-right UMAP displays the cell type assignment before reclustering. Therefore, during the annotation process, a reclustering on the NK and T cell was performed to gain in sub-population granularity and better discriminate CD8 / CD4 T cells. **e** The final population assignation.

### *Sig-annot*

#### Supplementary Figure S2

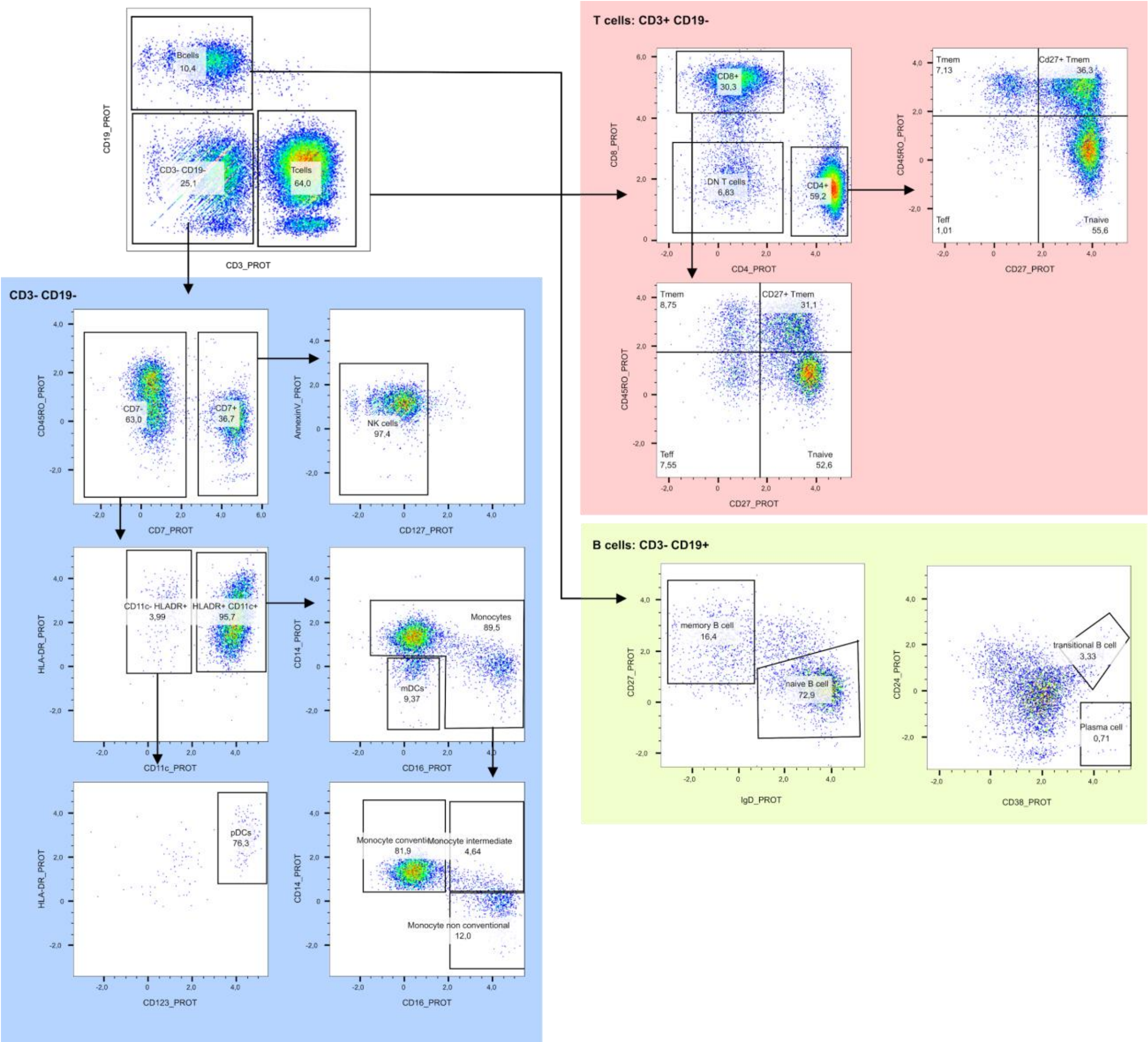

**Supplementary Figure S2** The gating strategy used to identify individual cell populations in the Kotliarov2020 dataset [1] based on the CITE-seq counts. The normalized protein counts were exported to FCS files using the R package flowCore [2,3] and loaded into FlowJo™ Software. Gating of individual cell populations was based on the gating strategy utilized in [4].

### Supplementary Figure S3

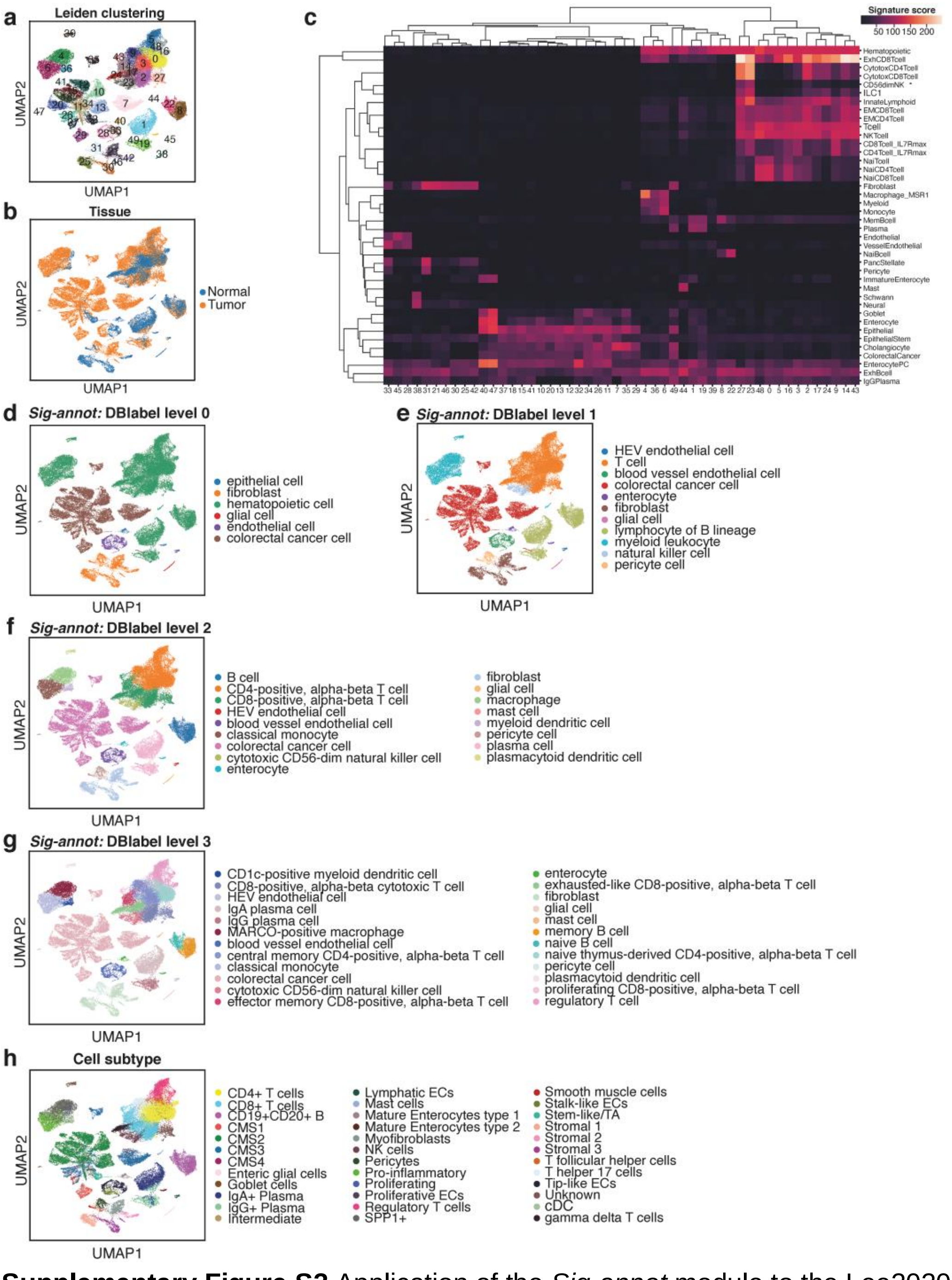

**Supplementary Figure S3** Application of the *Sig-annot* module to the Lee2020 dataset that includes tumor and non-malignant colon samples from colorectal cancer (CRC) patients [5]. **a** Leiden clustering. **b** Tumor and non-malignant normal control cell annotation. **c** Hierarchically clustered heatmap showing enrichment of main signatures employed in the annotation across leiden clusters, facilitating the evaluation of cluster attribution. **d** *Sig-annot* cell type attribution at level 0. **e** *Sig-annot* cell type attribution at level 1. **f** *Sig-annot* cell type attribution at level 2 **g** *Sig-annot* cell type attribution at level 3 **h** Author's cell type annotation at the highest resolution.

### Supplementary Figure S4

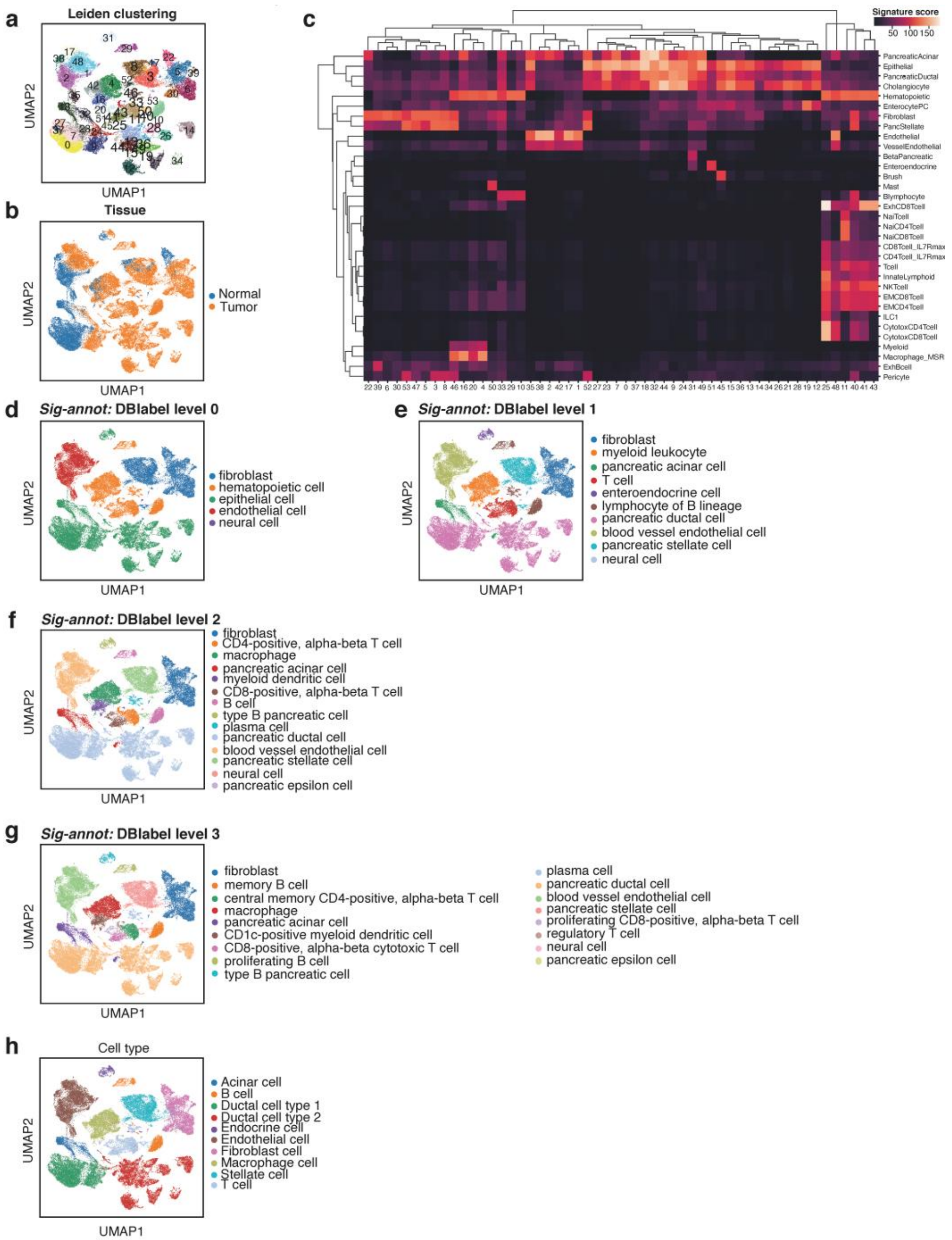

**Supplementary Figure S4** Application of the *Sig-annot* module to the Peng2019 dataset that includes tumor and non-malignant pancreatic samples from pancreatic ductal adenocarcinoma (PDAC) and non-pancreatic tumor patients [6]. **a** Leiden clustering. **b** Tumor and non-malignant norm al control cell annotation. **c** Hierarchically clustered heatmap showing enrichment of main signatures employed in the annotation across Leiden clusters, facilitating the evaluation of cluster attribution. **d** *Sig-annot* cell type attribution at level 0. **e** *Sig-annot* cell type attribution at level 1. **f** *Sig-annot* cell type attribution at level 2 **g** *Sig-annot* cell type attribution at level 3 **h** Author's cell type annotation.

### *Auto-annot*

#### Supplementary Figure S5

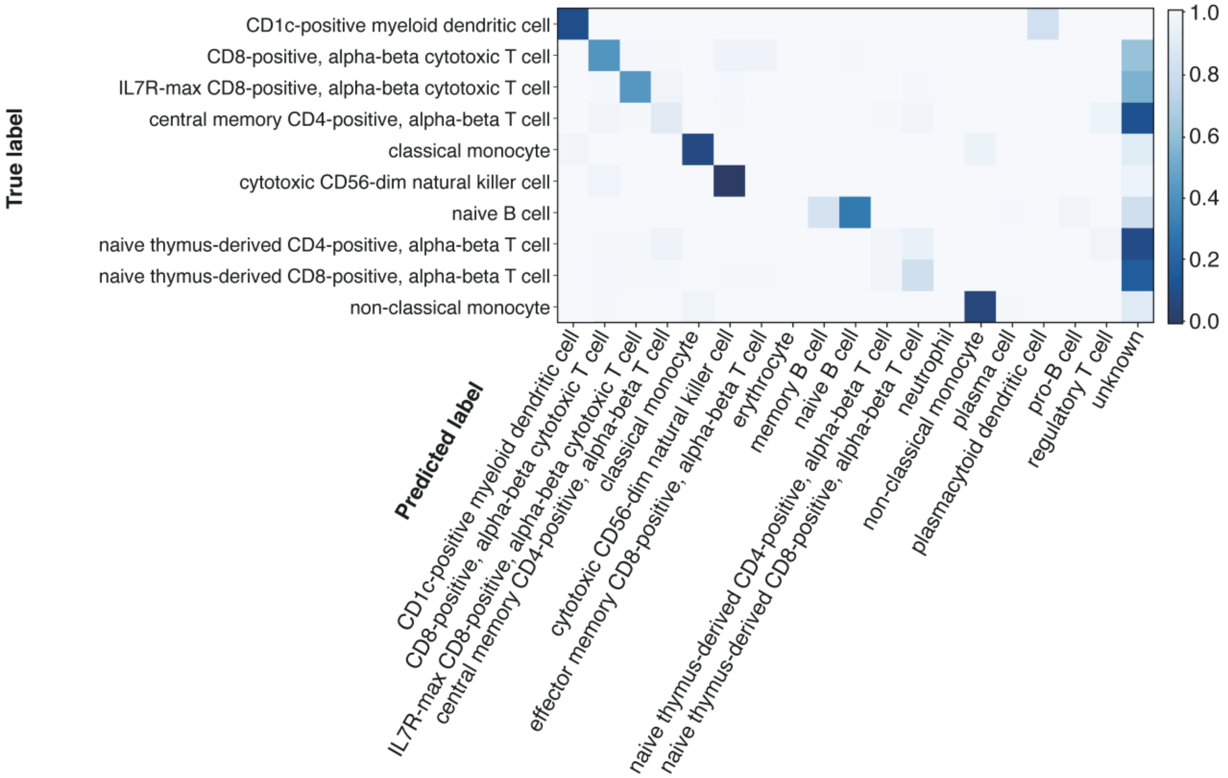

**Supplementary Figure S5** Application of the *Auto-annot* module to the Kotliarov2020 [1] and Granja2019 [7] as training data for a logistic regression model and the PBMC3k (https://www.10xgenomics.com/) dataset as a test dataset with 0.7 threshold. The setting of a threshold leads to a vanishing amount of misclassifications, at the cost of more unlabelled cells (“unknown”).

### Optimised Classes

Automated annotation based on less fine-grained cell types leads to clearer results (Supplementary Figure S6), as the problem that multiple different cell types being co-located in the same broad cell type class in another data set does not occur. In T cells some subtype borders are predicted differently by *Auto-annot*. This is likely to be due to the biological similarity and the smooth transition between cell types. Otherwise predictive performance is very high when the labels of the training and testing set are of identical annotation depth. Using a cut-off for prediction confidence scores highlights notoriously difficult to predict regions.

### Supplementary Figure S6

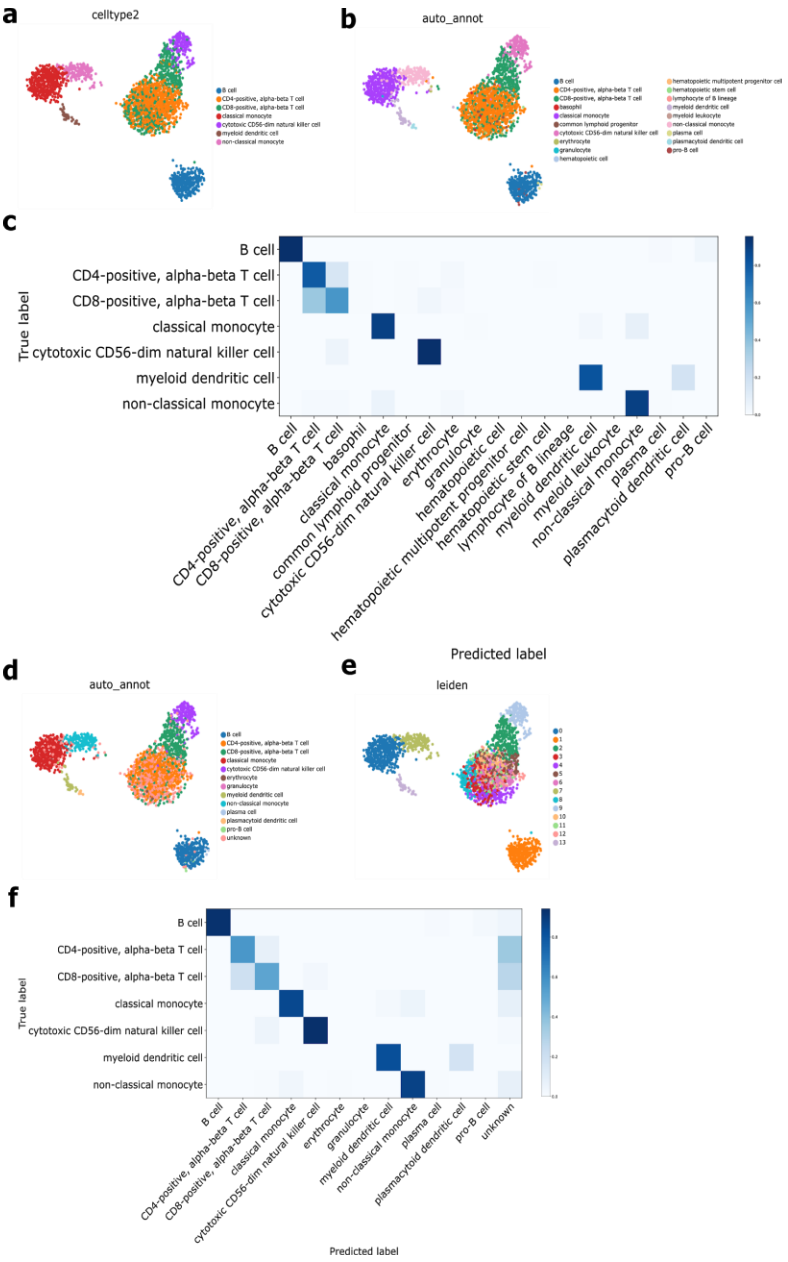

**Supplementary Figure S6** Optimised Classes. **a** Labels considered to be true in analysis. **b** Prediction based on the two training datasets Kotliarov2020 [1] and Granja2019 [7]. **c** Confusion matrix of that prediction. **d** Annotation where predictions with a low score are labelled as unknown. This removes many previously misclassified T cells and also misclassifications into cell types not present, which were already rare before. **e** The annotation shows a similar structure as Leiden clustering. **f** It can be seen that T cells on the fuzzy border between CD8 and CD4 are now more likely to be classified as unknown. If required, this behaviour can be tweaked by modifying the classification threshold.

### Supplementary Figure S7

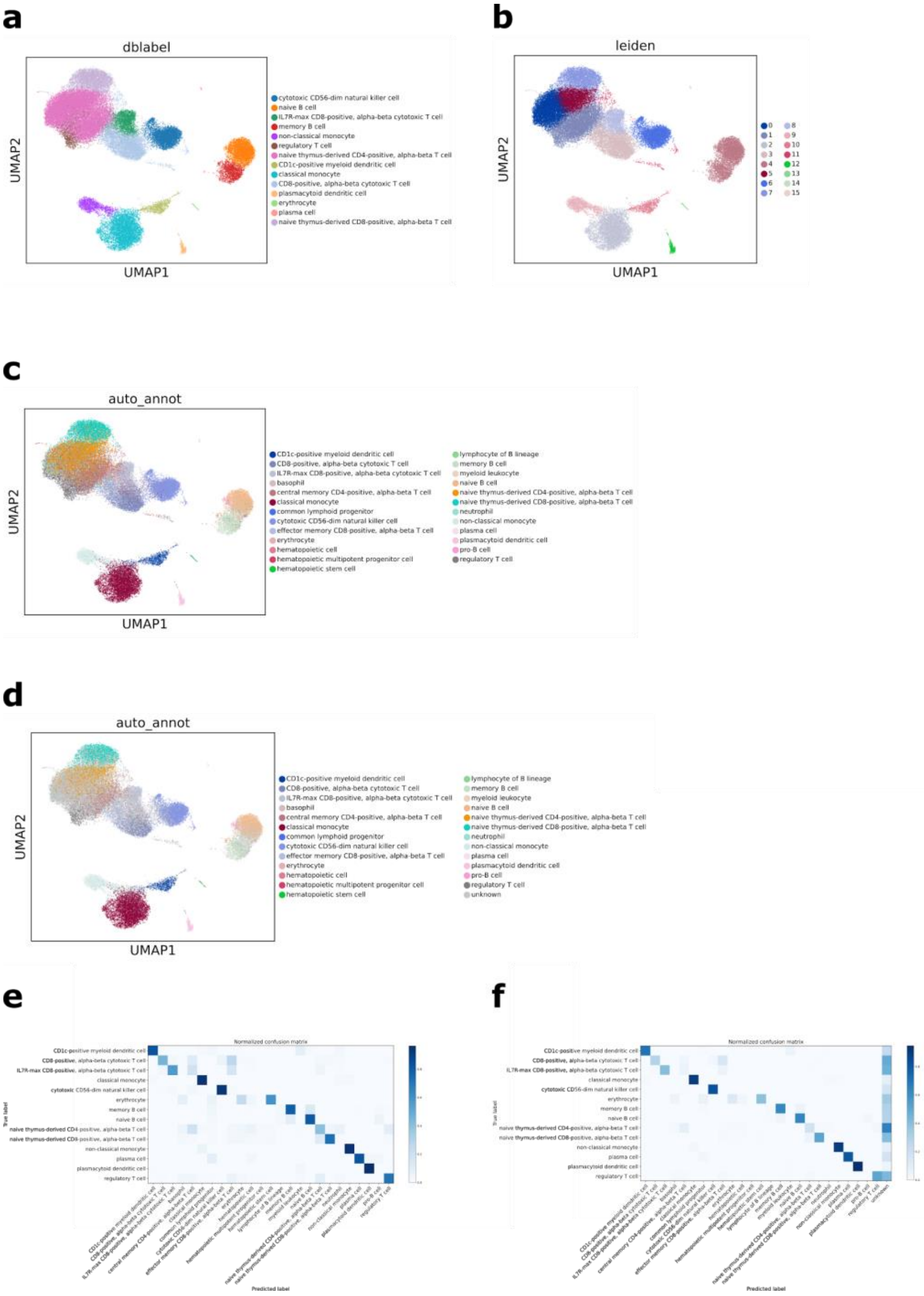

**Supplementary Figure S7** Train on Granja2019 [7], test in Kotliarov2020 [1]. **a** True labels from manual annotation. **b** Clusters generated from Leiden clustering. **c** Prediction through auto_annot reveals all major cell types with a finer graining of CD4 types and new hypotheses for small clusters. **d** Setting a confidence threshold (0.7) labels T cells whose subtype is unclear as unknown. **e** Confusion Matrix shows most of the misclassification happen in T cell subtypes. **f** Confusion matrix of prediction with confidence cut-off (0.7). The prediction accuracy seems greatly improved at the cost of retaining unlabelled cells.

### Supplementary Figure S8

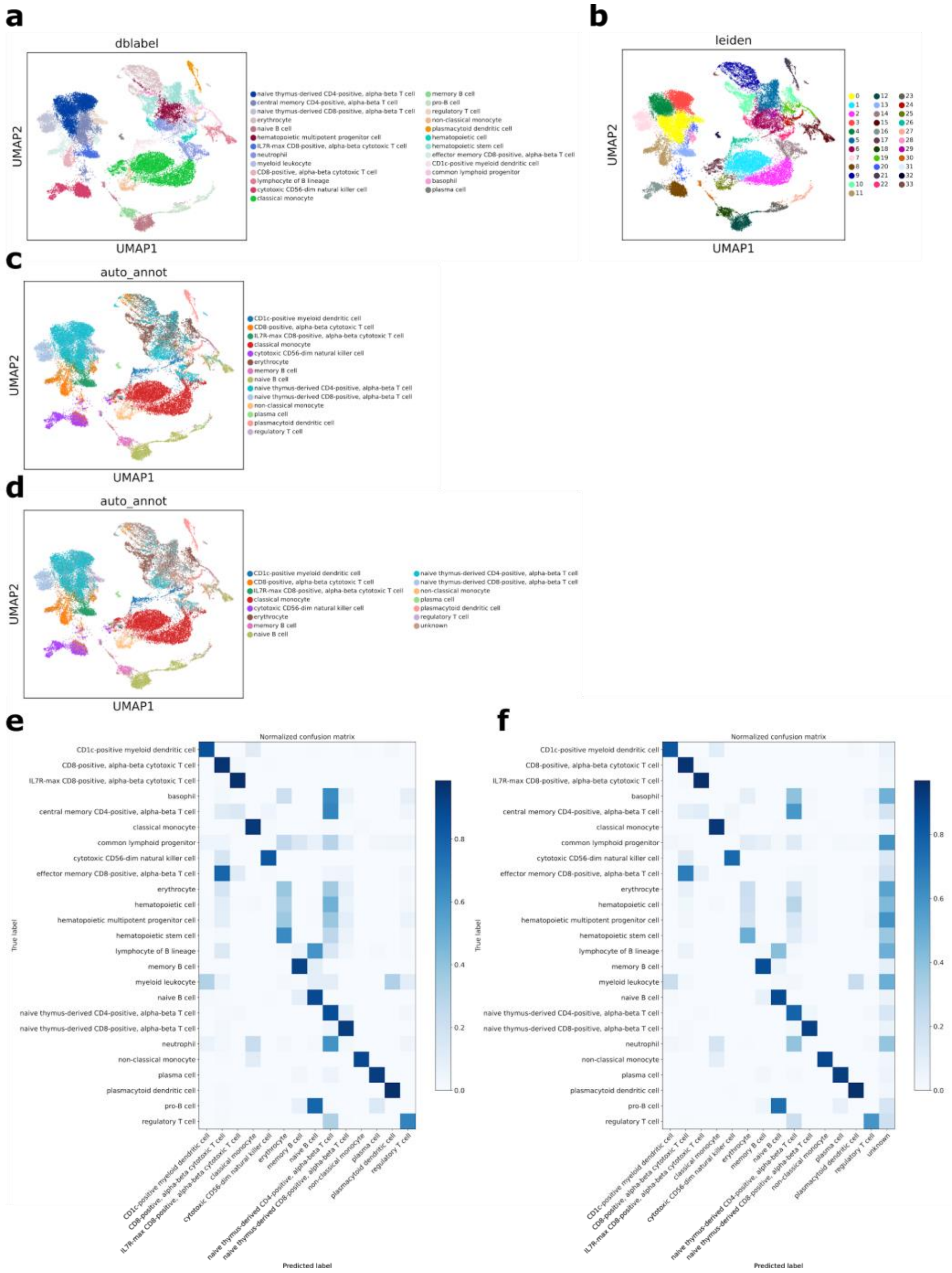

**Supplementary Figure S8** Train on Kotliarov2020 [1], test in Granja2019 [7]. **a** True labels from manual annotation. **b** Clusters generated from Leiden clustering. **c** The prediction gets all major cell types correct but cannot quite keep up with the fine grainedness of the testing set due to the limitations of the training set. **d** Some but not all of the misclassifications can be removed through setting a confidence cut-off (0.7). **e** The confusion matrix highlights difficulties in one to one comparison of predictions to truth when annotations are not identically structured and when some cell types are not present in both datasets. **f** Setting a cut-off (0.7) partially resolves this issue by allowing the classifier to only annotate cells where a certain confidence is reached.

### Supplementary Figure S9

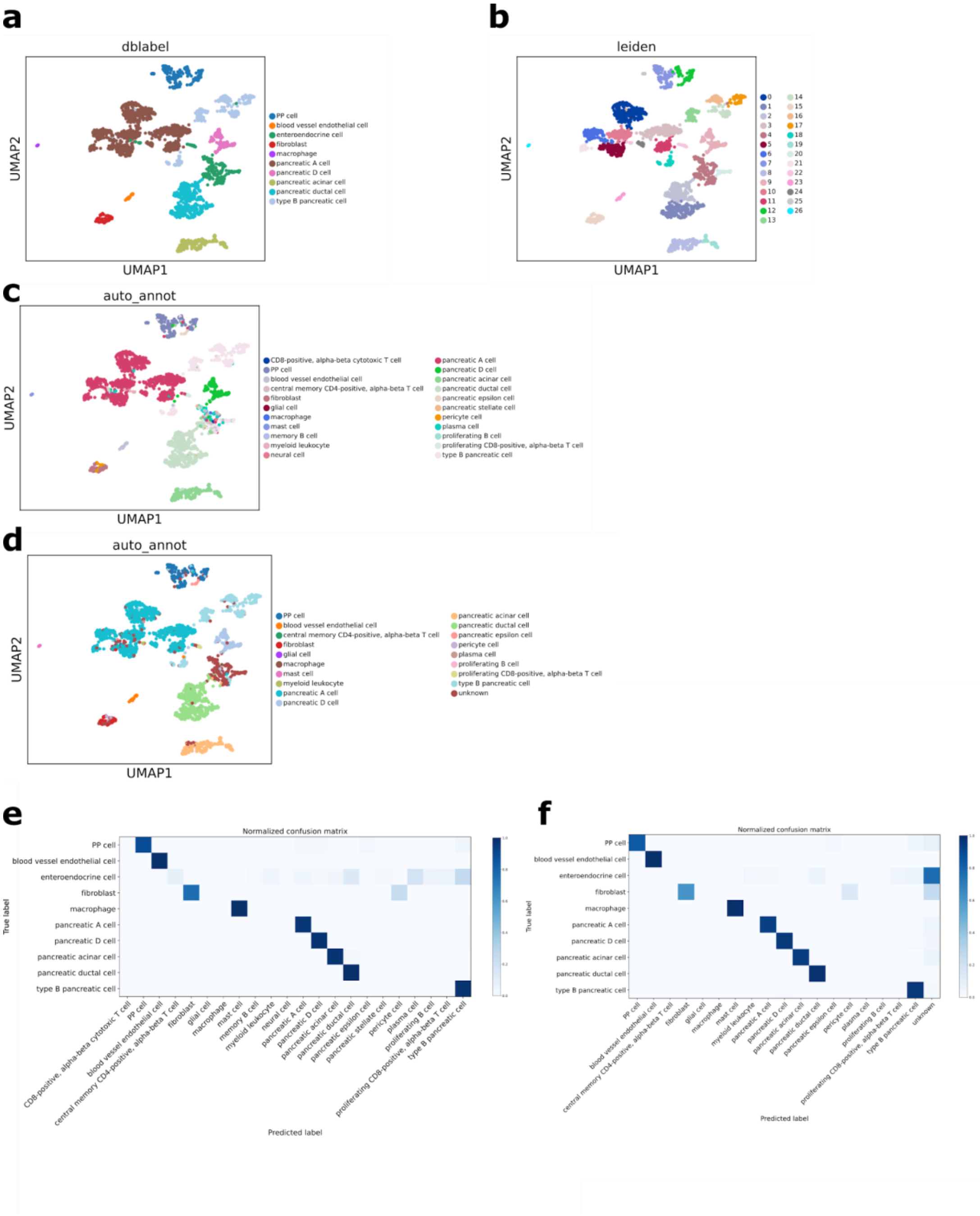

**Supplementary Figure S9** Train on Peng2019 [6] and Baron2016 [8], test in Segerstolpe2016 [9]. **a** True labels from manual annotation. **b** Clusters generated from Leiden clustering. **c** Most cells are labelled correctly, although individual cells may be misclassified. Enteroendocrine cells are always misclassified as they are not present in the training sets. **d** Using a confidence threshold (0.7) the enteroendocrine cell cluster can be labelled as unknown. **e** Most likely due to the small size of the testing set, annotation performance is excellent in all but enteroendocrine cells. **f** Setting a cut-off (0.7) avoids misclassification of enteroendocrines.

### Supplementary Figure S10

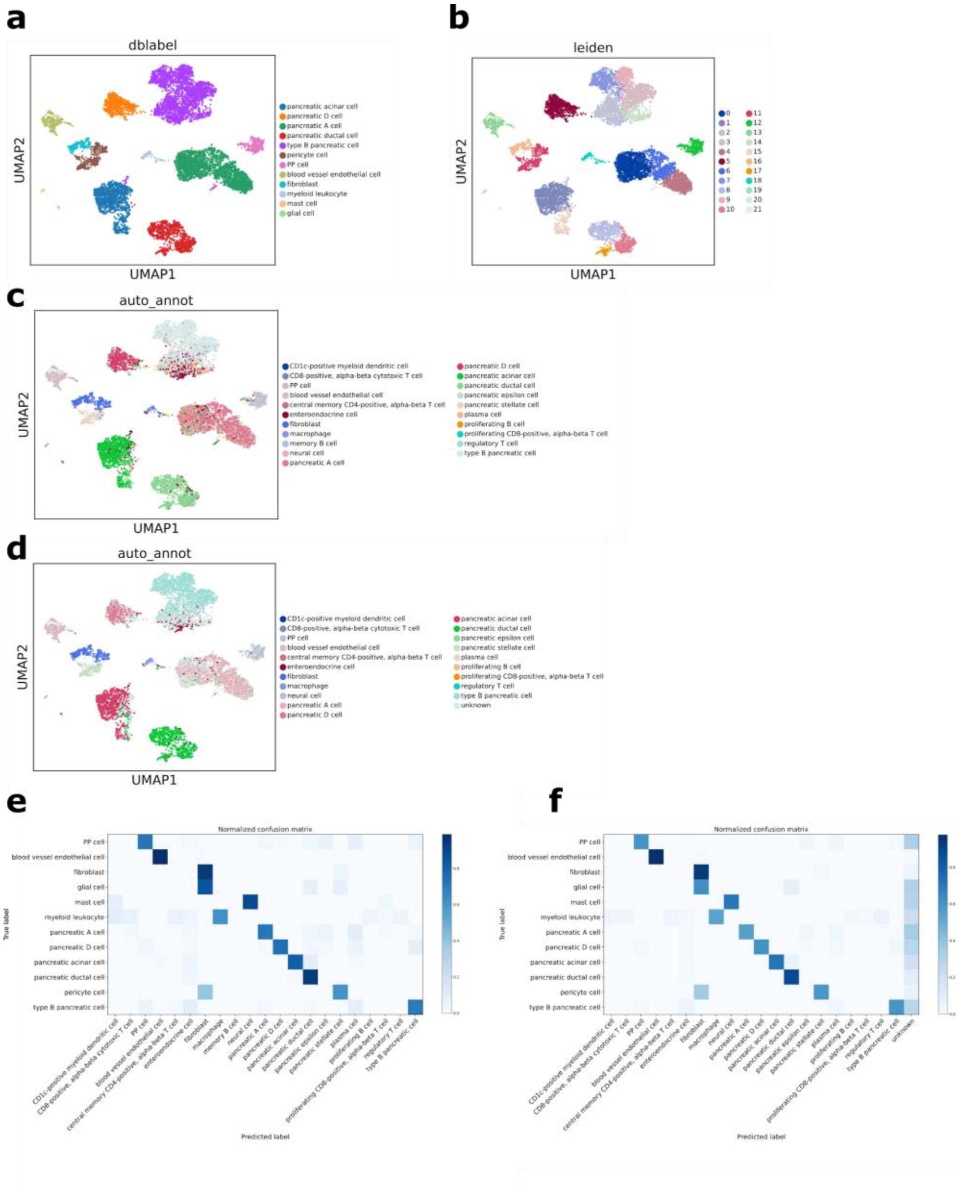

**Supplementary Figure S10** Train on Peng2019 [6] and Segerstolpe2016 [9], test in Baron2016 [8]. **a** True labels from manual annotation. **b** Clusters generated from Leiden clustering. **c** Generally, cells are labelled correctly, the inclusion of enteroendocrine cells from the Segerstolpe testing set, is in some parts erroneous, in other they may indicate a possible presence of such cells, and in all cases they elucidate the problem of unequal sets of cell types being present in training and testing. **d** A confidence cut-off (0.7) labels cells with a low confidence score as unknown, and further strengthens the hypothesis that a small enteroendocrine cluster may be present. **e** The confusion matrix shows that most types are correctly annotated, and also that the enteroendocrine cells originate from various cell types instead of all coming from the same true cell type label. **f** Using a confidence threshold (0.7) many of the previously misannotated cell types are annotated as unknown, but the effect does not affect a cell type in particular.

### Supplementary Figure S11

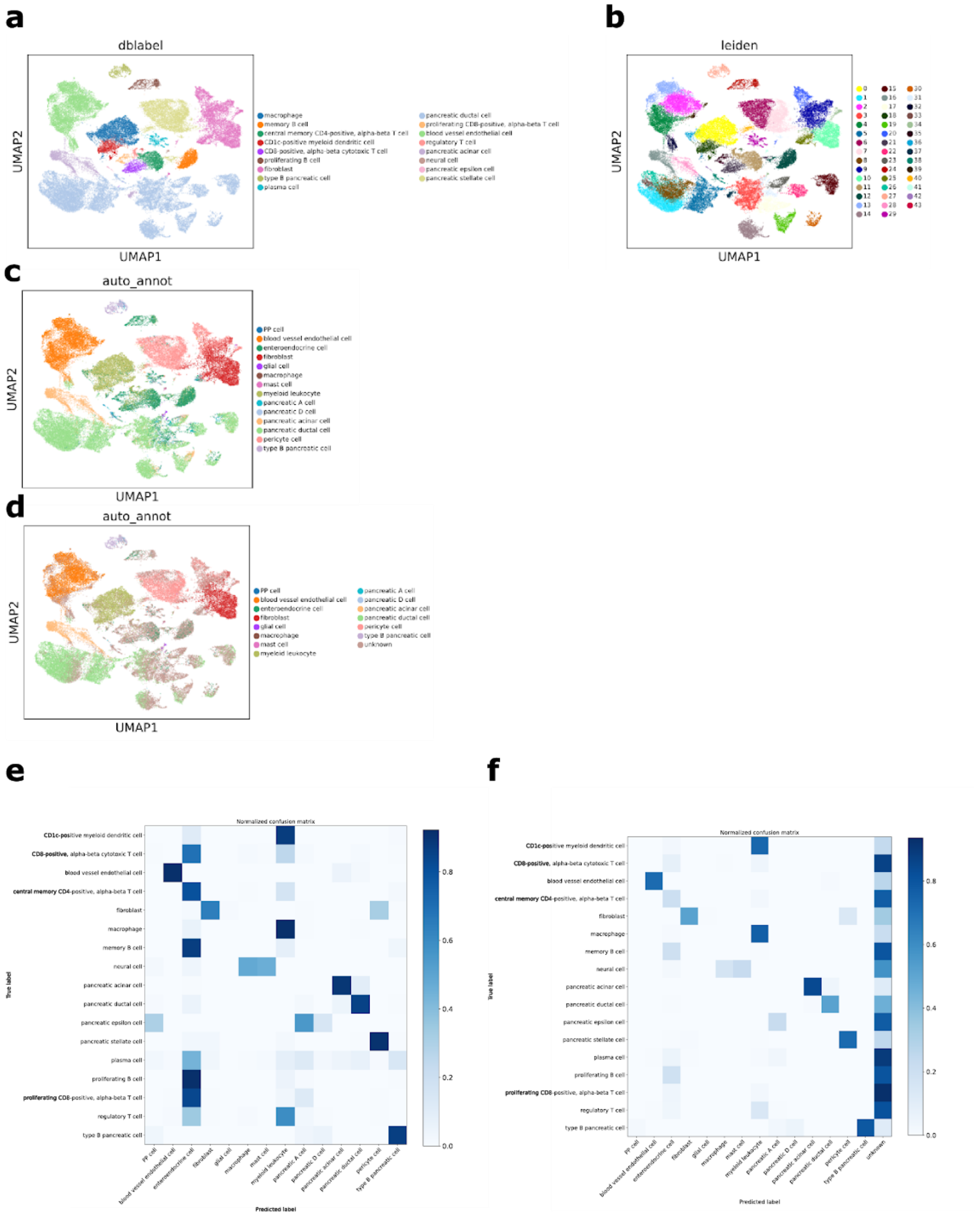

**Supplementary Figure S11** Train on Baron2016 [8] and Segestolpe2016 [9], test in Peng2019 [6]. **a** True labels from manual annotation. **b** Clusters generated from Leiden clustering. **c** *Auto-annot* correctly annotates most cell types but does not have sufficient granularity in the training dataset to replicate the fine grained original annotation. **d** The confidence threshold (0.7) removes many misclassified cells, but also removes some cells that were annotated correctly. **e** The confusion matrix highlights the difficulty of annotation when there is little overlap between the training classes and testing classes. **f** A confidence threshold (0.7) is able to circumvent this problem at the cost of leaving a large fraction of cells unlabelled.

### Supplementary Figure S12

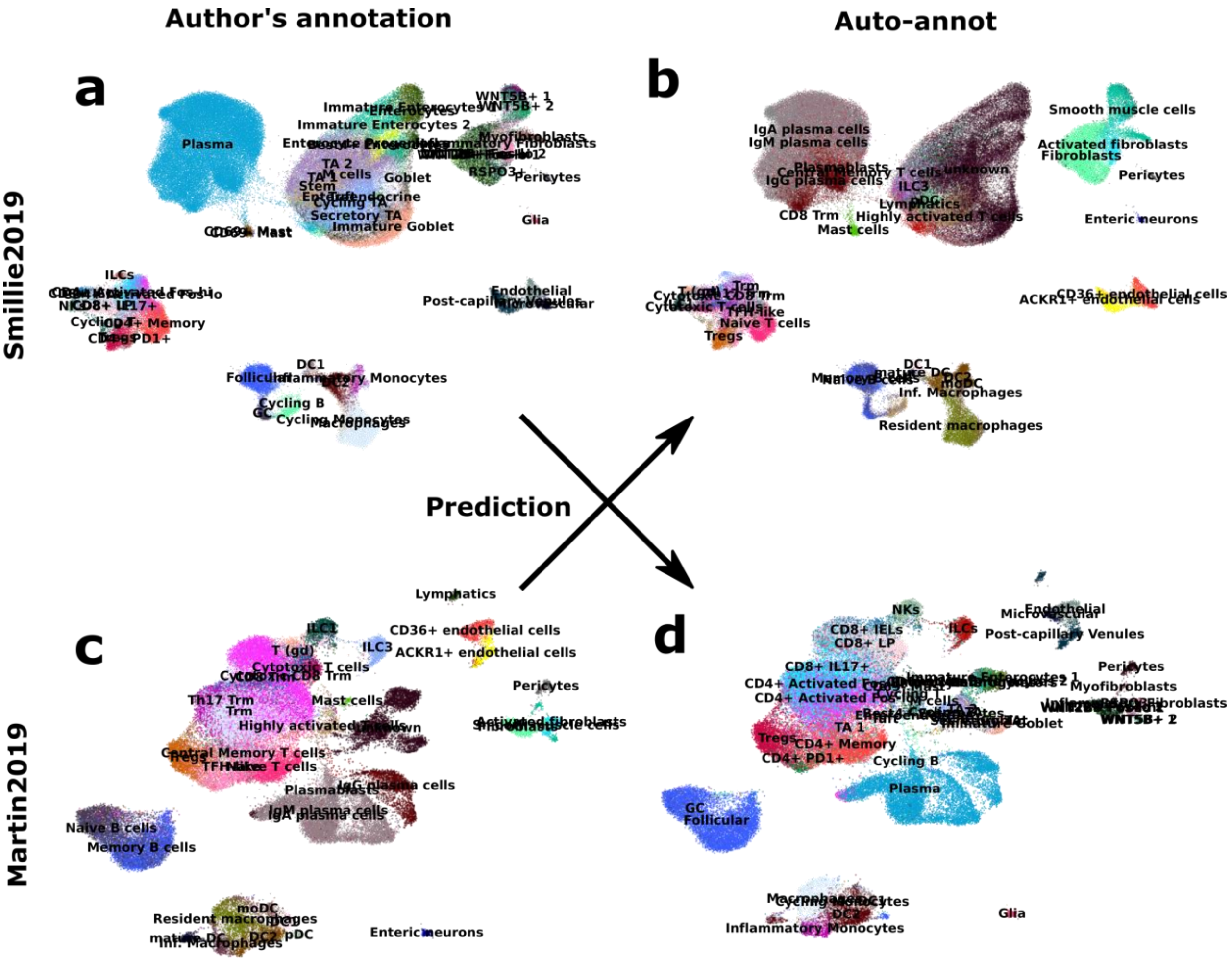

**Supplementary Figure S12** *Auto-annot* comparison between Smillie2019 [10] and Martin2019 [11]. **a-d** UMAP representations of the fine-grained cell types annotated in the Smillie2019 and Martin2019 datasets based on author's annotations (a,c) and predictions based on *Besca*'s *Auto-annot* module (b,d).

### Supplementary Figure S13

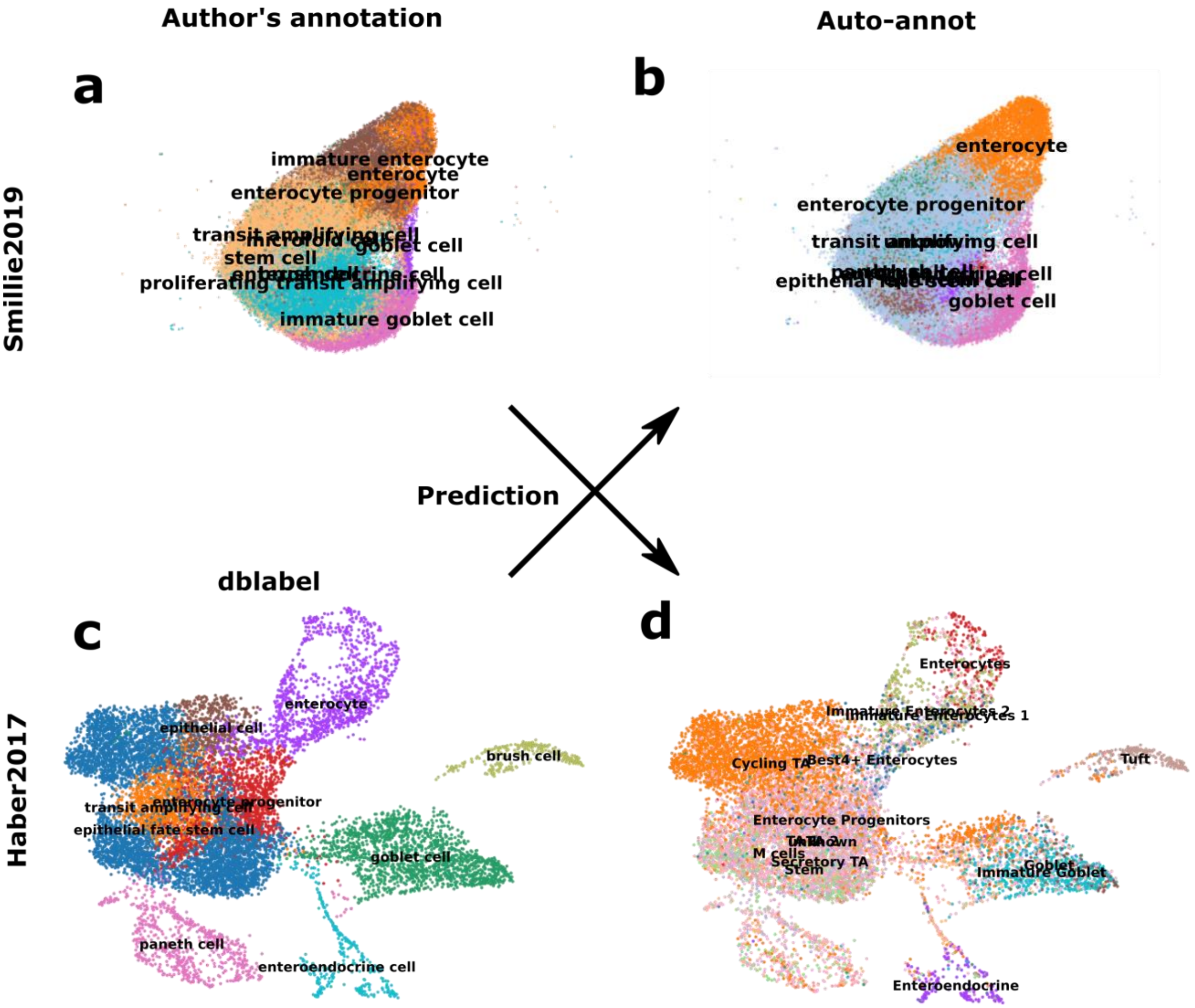

**Supplementary Figure S13** *Auto-annot* cross-species comparison between human Smillie2019 [10] and mouse Haber2017 [12]. **a-d** UMAP representations of the fine-grained author's annotation in the Smillie2019 dataset and DBlabel annotation in the Haber2017 dataset (a,c), and predictions based on *Besca*'s *Auto-annot* module (b,d).

### *Bescape*

#### Supplementary Figure S14

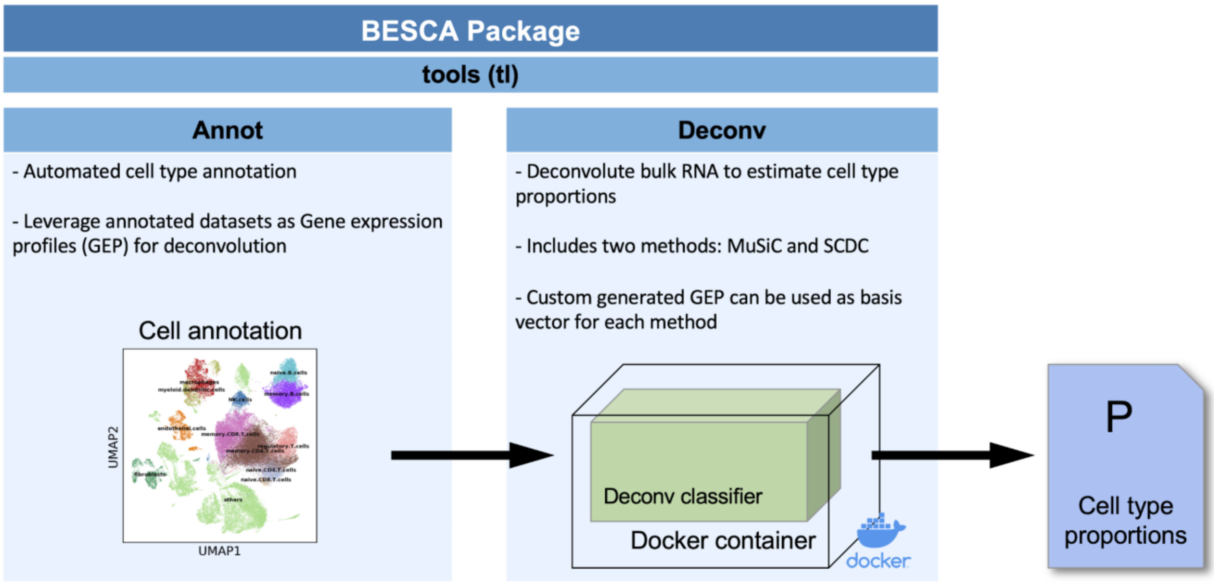

**Supplementary Figure S14** *Bescape*, a containerized environment for cell deconvolution based on *Besca*'s annotation modules.

### Supplementary Figure S15

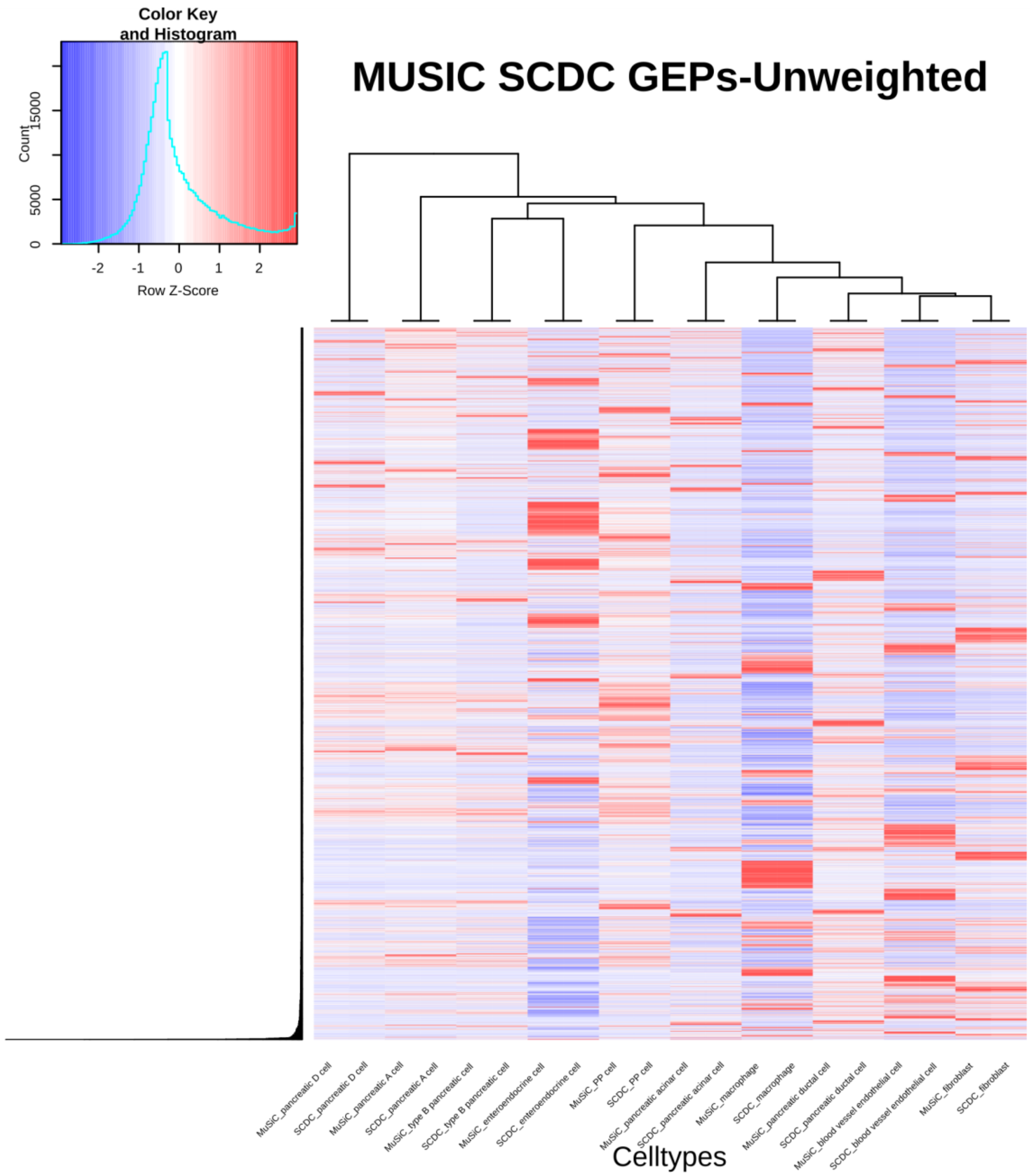

**Supplementary Figure S15** Heatmap of the gene expression profile (GEP) generated by the basis matrix generation as part of SCDC [13] and MuSiC [14] methods taking in directly the raw counts of the pancreatic reference scRNA-seq dataset [9] and applying weights to all 21246 different genes without any feature selection needed.

### Supplementary Figure S16

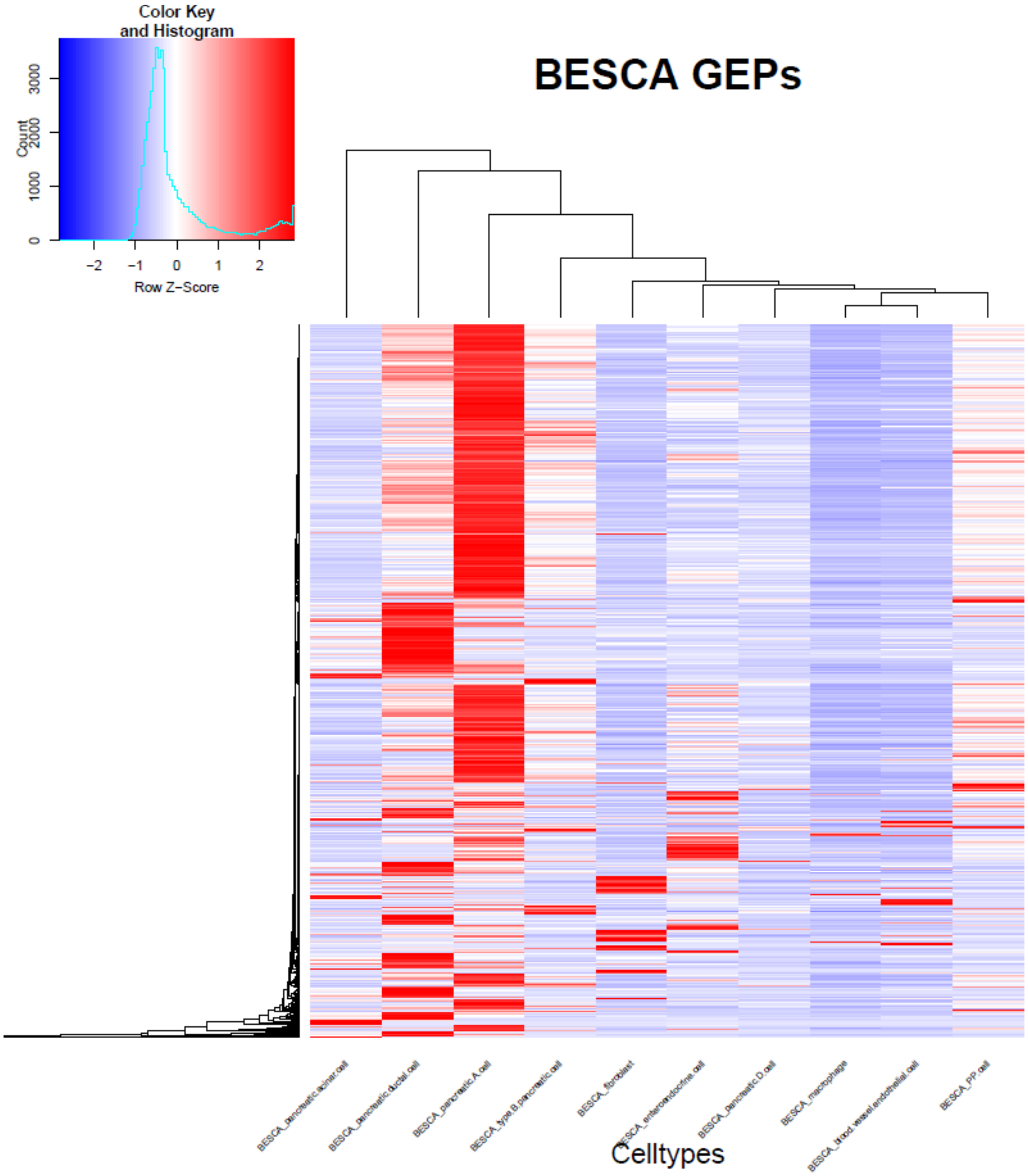

**Supplementary Figure S16** Heatmap of the gene expression profile (GEP) generated by the output of the *Besca* workflow from selection of genes based on known markers and level of variability seen across annotated cell types, resulting in some 5140 informative genes in the pancreatic reference scRNA-seq dataset [9].

### Supplementary Figure S17

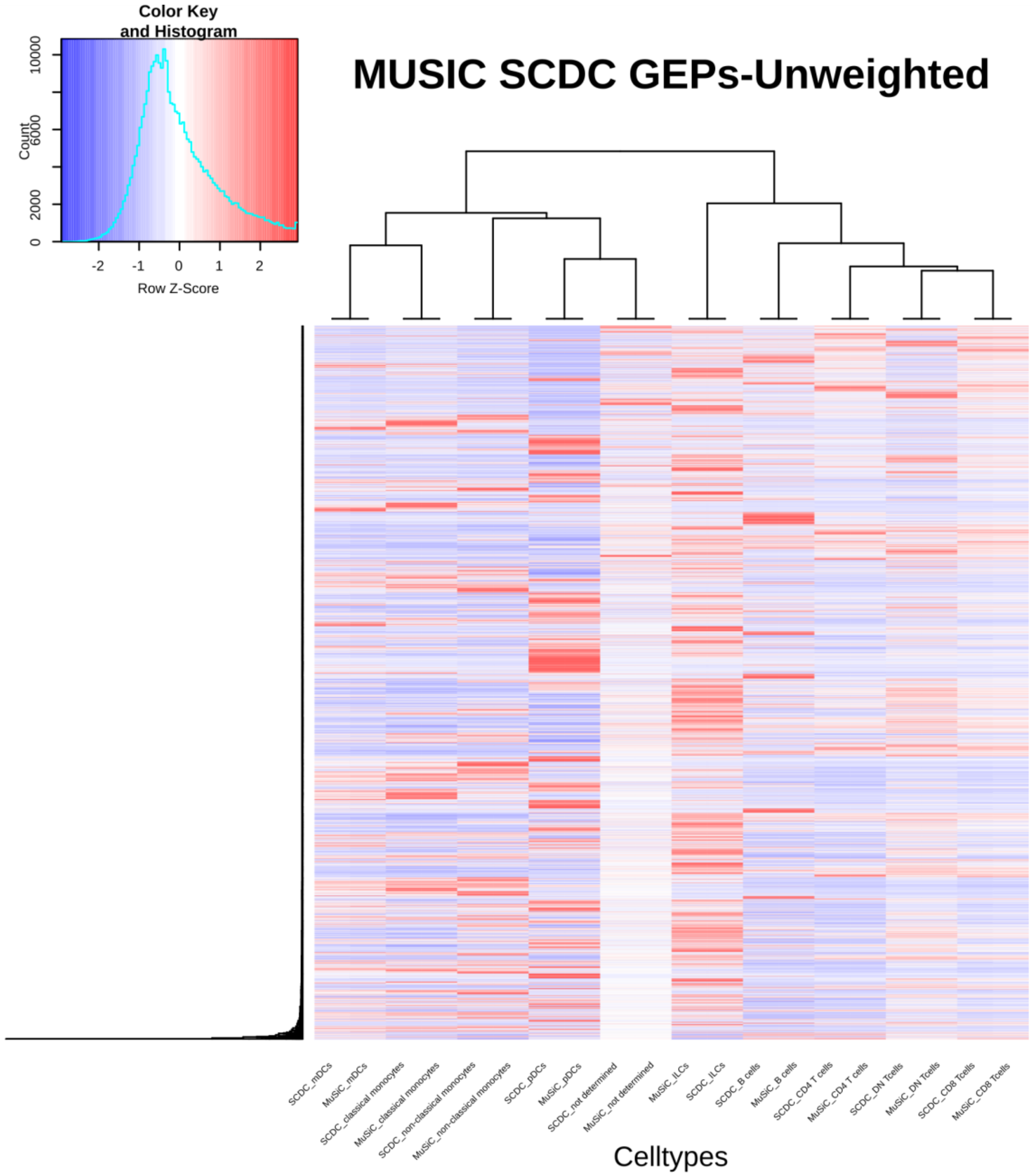

**Supplementary Figure S17** Heatmap of the gene expression profile (GEP) generated by the basis matrix generation as part of SCDC [13] and MuSiC [14] methods taking in directly the raw counts of the hematopoietic reference CITE-seq dataset [1] and applying weights to all 14935 different genes without any feature selection needed.

### Supplementary Figure S18

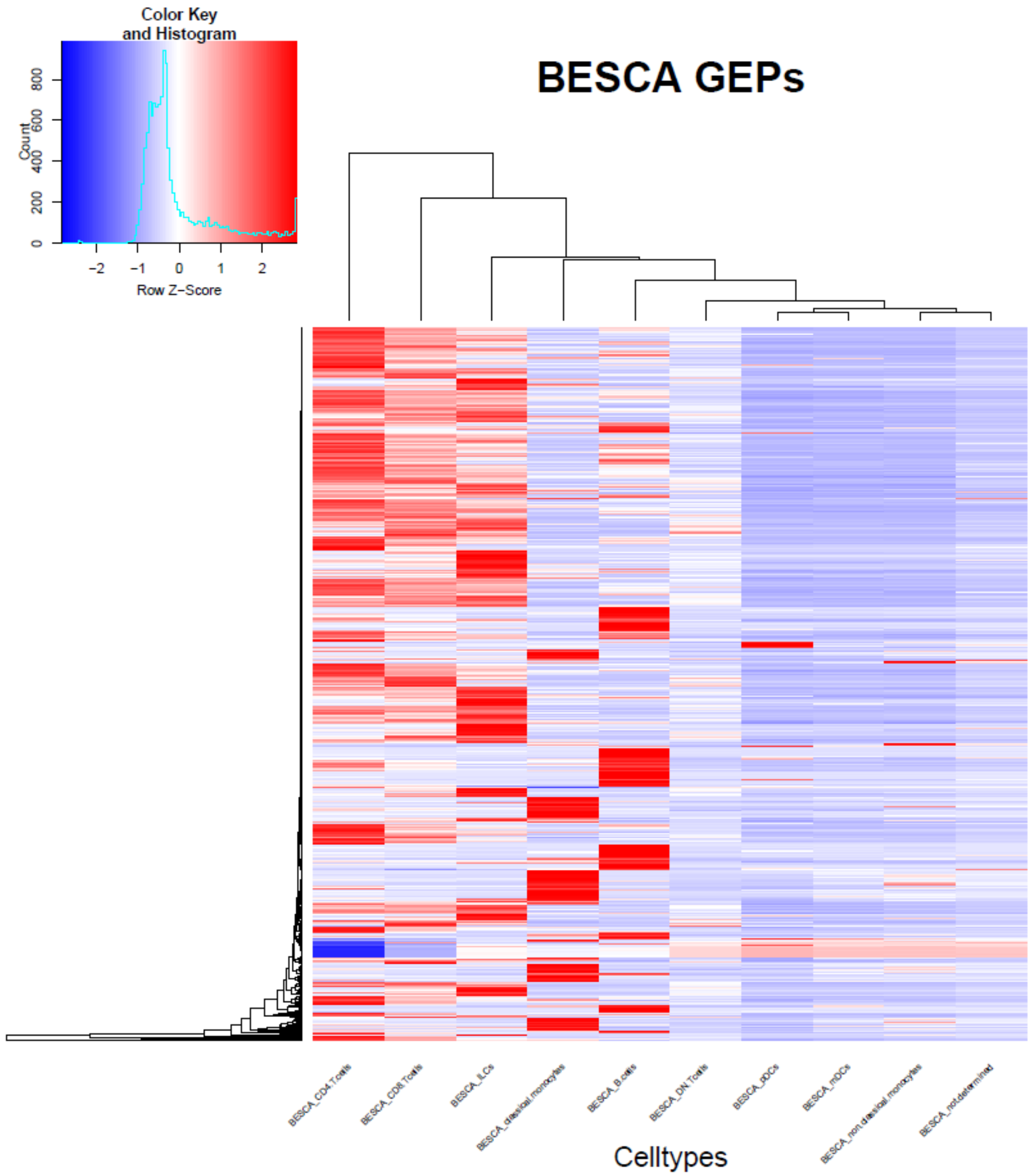

**Supplementary Figure S18** Heatmap of the gene expression profile (GEP) generated by the output of the *Besca* workflow from selection of genes based on known markers and level of variability seen across annotated cell types, resulting in some 1271 informative genes in the hematopoietic reference CITE-seq dataset [1].

### References

1. Kotliarov Y, Sparks R, Martins AJ, Mulè MP, Lu Y, Goswami M, et al. Broad immune activation underlies shared set point signatures for vaccine responsiveness in healthy individuals and disease activity in patients with lupus. Nat Med. Nature Publishing Group; 2020;26:618–29.

2. Hahne F, LeMeur N, Brinkman RR, Ellis B, Haaland P, Sarkar D, et al. flowCore: a Bioconductor package for high throughput flow cytometry. BMC Bioinformatics. 2009;10:106.

3. Ellis B, Haal P, Hahne F, Meur NL, Gopalakrishnan N, Spidlen J, et al. flowCore: flowCore: Basic structures for flow cytometry data [Internet]. Bioconductor version: Release (3.11); 2020 [cited 2020 Aug 2]. Available from: https://bioconductor.org/packages/flowCore/

4. Waugh KA, Araya P, Pandey A, Jordan KR, Smith KP, Granrath RE, et al. Mass Cytometry Reveals Global Immune Remodeling with Multi-lineage Hypersensitivity to Type I Interferon in Down Syndrome. Cell Rep. 2019;29:1893-1908.e4.

5. Lee H-O, Hong Y, Etlioglu HE, Cho YB, Pomella V, Van den Bosch B, et al. Lineage-dependent gene expression programs influence the immune landscape of colorectal cancer. Nat Genet. Nature Publishing Group; 2020;52:594–603.

6. Peng J, Sun B-F, Chen C-Y, Zhou J-Y, Chen Y-S, Chen H, et al. Single-cell RNA-seq highlights intra-tumoral heterogeneity and malignant progression in pancreatic ductal adenocarcinoma. Cell Res. Nature Publishing Group; 2019;29:725–38.

7. Granja JM, Klemm S, McGinnis LM, Kathiria AS, Mezger A, Corces MR, et al. Single-cell multiomic analysis identifies regulatory programs in mixed-phenotype acute leukemia. Nat Biotechnol. Nature Publishing Group; 2019;37:1458–65.

8. Baron M, Veres A, Wolock SL, Faust AL, Gaujoux R, Vetere A, et al. A Single-Cell Transcriptomic Map of the Human and Mouse Pancreas Reveals Inter- and Intra-cell Population Structure. Cell Syst. Elsevier; 2016;3:346-360.e4.

9. Segerstolpe Å, Palasantza A, Eliasson P, Andersson E-M, Andréasson A-C, Sun X, et al. Single-Cell Transcriptome Profiling of Human Pancreatic Islets in Health and Type 2 Diabetes. Cell Metab. Elsevier; 2016;24:593–607.

10. Smillie CS, Biton M, Ordovas-Montanes J, Sullivan KM, Burgin G, Graham DB, et al. Intra- and Inter-cellular Rewiring of the Human Colon during Ulcerative Colitis. Cell. Elsevier; 2019;178:714-730.e22.

11. Martin JC, Chang C, Boschetti G, Ungaro R, Giri M, Grout JA, et al. Single-Cell Analysis of Crohn’s Disease Lesions Identifies a Pathogenic Cellular Module Associated with Resistance to Anti-TNF Therapy. Cell. Elsevier; 2019;178:1493-1508.e20.

12. Haber AL, Biton M, Rogel N, Herbst RH, Shekhar K, Smillie C, et al. A single-cell survey of the small intestinal epithelium. Nature. Nature Publishing Group; 2017;551:333–9.

13. Dong M, Thennavan A, Urrutia E, Li Y, Perou CM, Zou F, et al. SCDC: bulk gene expression deconvolution by multiple single-cell RNA sequencing references. Brief Bioinform. 2020;

14. Wang X, Park J, Susztak K, Zhang NR, Li M. Bulk tissue cell type deconvolution with multi-subject single-cell expression reference. Nat Commun. Nature Publishing Group; 2019;10:380.
